# IPMK regulates HDAC3 activity and histone H4 acetylation in human cells

**DOI:** 10.1101/2024.04.29.591660

**Authors:** Gregory A. Sowd, Elizabeth A. Stivison, Pratima Chapagain, Andrew T. Hale, James C. Poland, Lucia E. Rameh, Raymond D. Blind

## Abstract

Histone deacetylases (HDACs) repress transcription by catalyzing the removal of acetyl groups from histones. Class 1 HDACs are activated by inositol phosphate signaling molecules *in vitro*, but it is unclear if this regulation occurs in human cells. Inositol Polyphosphate Multikinase (IPMK) is required for production of inositol hexakisphosphate (IP6), pentakisphosphate (IP5) and certain tetrakisphosphate (IP4) species, all known activators of Class 1 HDACs *in vitro*. Here, we generated IPMK knockout (IKO) human U251 glioblastoma cells, which decreased cellular inositol phosphate levels and increased histone H4-acetylation by mass spectrometry. ChIP-seq showed IKO increased H4-acetylation at IKO-upregulated genes, but H4-acetylation was unchanged at IKO-downregulated genes, suggesting gene-specific responses to IPMK knockout. HDAC deacetylase enzyme activity was decreased in HDAC3 immunoprecipitates from IKO *vs*. wild-type cells, while deacetylase activity of other Class 1 HDACs had no detectable changes in activity. Wild-type IPMK expression in IKO cells fully rescued HDAC3 deacetylase activity, while kinase-dead IPMK expression had no effect. Further, the deficiency in HDAC3 activity in immunoprecipitates from IKO cells could be fully rescued by addition of synthesized IP4 (Ins(1,4,5,6)P4) to the enzyme assay, while control inositol had no effect. These data suggest that cellular IPMK-dependent inositol phosphates are required for full HDAC3 enzyme activity and proper histone H4-acetylation. Implications for targeting IPMK in HDAC3-dependent diseases are discussed.

## INTRODUCTION

Gene expression is regulated by the dynamic acetylation and de-acetylation of histones, which influence chromatin packing and other aspects of transcriptional regulation. Increased acetylation generally leads to increased chromatin accessibility and activated transcription, while decreased acetylation generally represses transcription^1–4^. Histone acetylation is controlled by the opposing activities of many enzymes, one group are histone deacetylases (HDACs)^5^, which function to enzymatically remove acetyl groups from histones and other proteins. While the molecular details of HDAC regulation remain incompletely understood, it is well established that purified, recombinant Class 1 HDACs are activated by inositol phosphates *in vitro*^6,7^. Inositol phosphates (IPs) are metabolites^8–10^ that bind function mutations only affect inositol phosphate binding. Gut microbiota derived inositol phosphates have been suggested to directly activate HDAC3 in the intestinal epithelium of mice^14^ and genetic studies with the yeast directly to a conserved IP-binding site close to the Class 1 HDACs active site to increase deacetylase activity^6,7^. Class 1 HDACs were discovered by crystallography to interact with the inositol phosphate Ins(1,4,5,6)P4, which fortuitously co-purified with the protein in the crystal structure of HDAC3^11^. Mutation of the conserved IP-binding residues abolished HDAC3 enzyme activity *in vitro*^11^ and deceased HDAC1/2 activity in living cells^12^. All Class I HDACs tested have activated enzyme kinetics in the presence of higher-order phosphorylated species of inositol phosphates^6,7,13^. However, the extent to which Class 1 HDACs require inositol phosphates *in vivo* remains unclear. Triple mutations in the inositol binding site change HDAC1/2-dependent phenotypes in mouse ES cells, but it is unclear if loss of orthologue of Class 1 HDACs, Rpd3^15^ demonstrate a clear connection between inositol phosphate synthesis genes and Rpd3 function in yeast^16^.

Although the connection between inositol phosphates and Class 1 HDAC activity is well supported, it still remains unknown if altering inositol phosphate levels in human cells alters histone acetylation, Class 1 HDAC enzyme activity and gene expression. Here, we attempt to address these questions using genetic deletion of Inositol Polyphosphate Multikinase (*IPMK*), an enzyme central to all higher order inositol phosphate biosynthesis in human cells.

Inositol Polyphosphate Multikinase (IPMK) is an inositol kinase conserved across all eukaryotes, operating at the nexus of several complex inositol phosphate signaling pathways^17–19^. IPMK catalyzes phosphorylation of IP3 to IP4, and IP4 to IP5. Thus, inositol pentakisphosphate (IP5) and inositol hexakisphosphate (IP6) are depleted from cells lacking IPMK^20^, while certain IP4 isoforms are synthesized independently of IPMK, by IP3K and ITPK1^21,22^. IPMK was first discovered as part of the MCM1 transcriptional regulatory complex^23–26^ and was only later identified to have inositol kinase activity^27^. IPMK regulates gene expression in both kinase-independent ^28^ and kinase-dependent ^29,30^ manners, however only a handful of studies have revealed mechanistic details of how IPMK kinase activity regulates transcription at the molecular level^31–33^.

Here, we used genetic deletion of *IPMK* to test if Class I HDAC function is affected in human cells by depletion of IP5 and IP6. *IPMK* deletion depleted all detectable IP5 and IP6, and upregulated histone H4 acetylation by mass spectrometry and ChIP-seq in human U251-MG cells, and the IPMK-regulated transcriptome was enriched in HDAC-regulated genes. The kinase activity of IPMK was required for full deacetylase activity of HDAC3 immunopurified from these cells, as expression of wild-type full restored HDAC3 activity, while kinase-dead IPMK expression had no effect. Addition of the IPMK-generated inositol phosphate enzyme product (Ins(1,4,5,6)P4) to the HDAC3 enzyme reaction completely rescued the deacetylase activity, while addition of control inositol had no effect. These data suggest IPMK kinase-dependent inositol phosphates are critical for full HDAC3 activity in human U251 cells, which can impact development of new therapies targeting HDAC3, as well as our basic understanding of how IPMK and inositol phosphates regulate transcription.

## RESULTS

### IPMK knockout decreases human cell growth and inositol phosphate levels

Inositol hexakisphosphate (IP6) was previously co-crystalized with HDAC1^6^ (**Fig 1A**) and activated the deacetylase activity of all class 1 HDACs tested *in vitro*^6,7^. Since IP6 and all higher-order inositol phosphate production in human cells depends on Inositol Polyphosphate Multikinase (IPMK, **Fig 1B**), we deleted *IPMK* from human U251-MG cells to investigate the effects of inositol phosphate depletion on HDAC activity *in vivo.* CRISPR-Cas9-based deletion of all *IPMK* alleles (IKO, **Fig 1C, Fig S1A-D**) depleted the expected inositol phosphate species in *IPMK* knockout (IKO) cells as measured by HPLC of cells metabolically radiolabeled with ^3^H-inositol (**Fig 1D**). Consistent with previous studies, IP5 and IP6 were undetectable upon IKO^34^. IP4 persisted which is likely an isoform of IP4 (Ins(1,3,4,5)P4) produced by IPMK-independent pathways^35^. IKO clones grew more slowly than on gene expression, we analyzed the transcriptomes of IKO relative to WT cells by RNA-seq, which showed differential expression of 4,418 genes (*p_adj_*<0.05) or 896 genes (*p_adj_*<0.05 and Log_2_FC±1.0) (**Fig 1F** and **Spreadsheet 1**). These IPMK-regulated transcripts were biased toward upregulation of transcript abundance in IKO cells (**Fig 1G**), consistent with IPMK functioning to actively repress gene expression. Indeed, enrichment of Class 1 HDAC gene sets were discovered by gene set enrichment analysis (**Fig 1H**, **Spreadsheet 2**). These data suggest that loss of IPMK prevents formation of higher order inositol phosphates, slows U251 human cell growth, and alters expression of genes known to be regulated by Class I HDACs.

**Figure 1.**
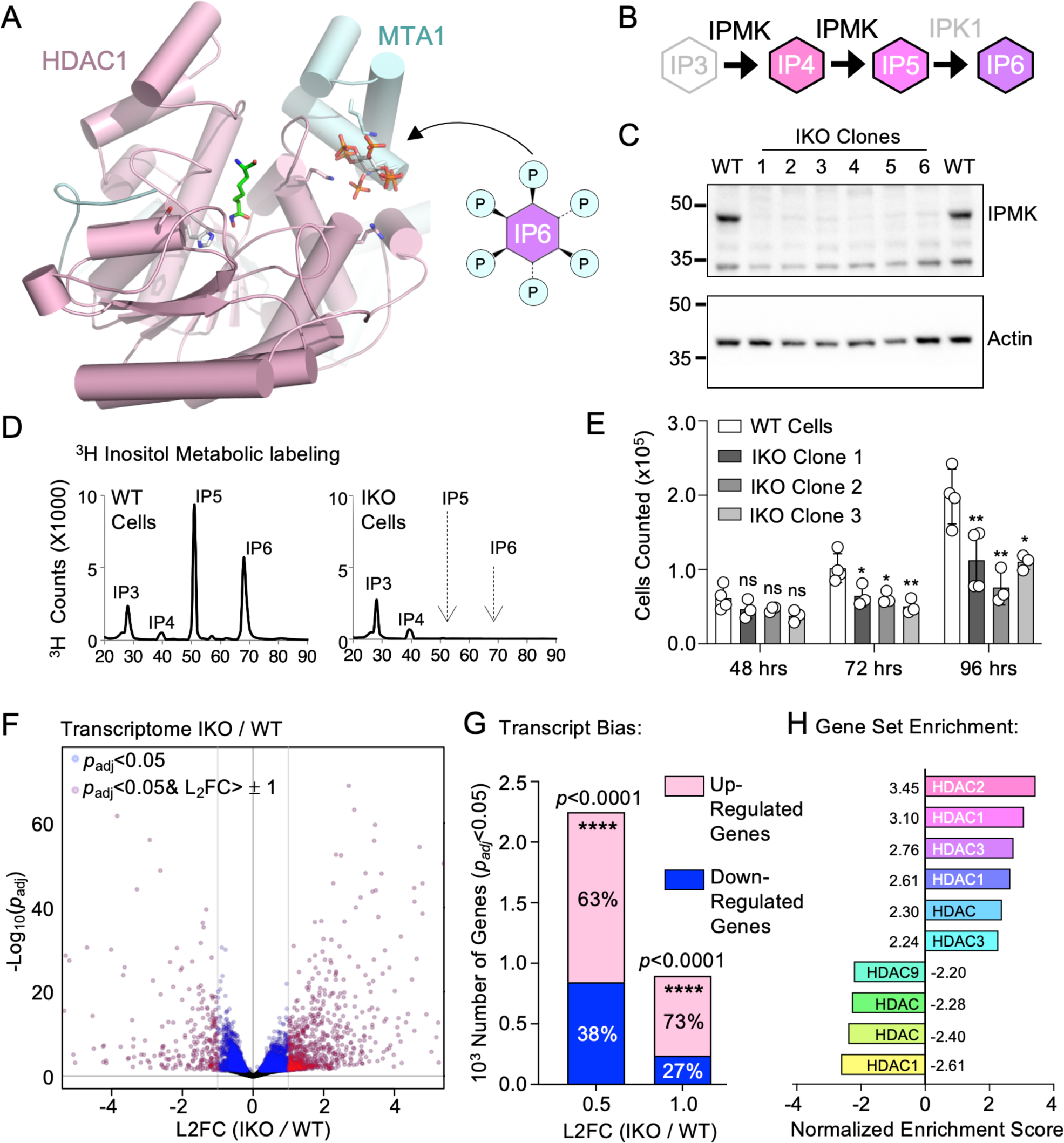
IPMK genetic deletion in human U251 cells upregulates gene expression, with high enrichment in HDAC gene sets. **A.** Published crystal structure of IP6 bound to HDAC1 (PDB:5ICN), illustrating the position of IP6 in the structure previously shown to increase HDAC enzyme activity. **B.** Role of IPMK in inositol phosphate metabolism, more detail provided in Figure 6. **C**. Western blot of WT *vs*. IPMK-knockout (IKO) human U251-MG whole cell lysates from six independent, monoclonal CRISPR clones, genomic DNA PCR showing IPMK knockout provided in supplemental. **D**. HPLC profiles of metabolically ^3^H-inositol-labeled WT (left) or IKO (right) U251 cells, migration of inositol phosphate standards indicated. **E**. Cell counts of wild-type vs. three independent monoclonal IKO CRISPR-clones counted after indicated times post-plating, \**p_adj_*<0.05, \*\**p_adj_*<0.01, in ordinary one-way ANOVA Dunnett’s corrected; ns is not significant. **F**. Volcano plot of RNA-seq from IKO *vs*. WT cells and **G**. contingency analysis of indicated genes, analyzed by Fisher’s exact test. **H**. Normalized gene set enrichment scores of HDAC gene sets from the IKO-regulated transcriptome, full sheets for GSEA supplied as supplemental. These data suggest genetic removal of IPMK decreases inositol phosphate levels and biases gene expression to be upregulated, with high enrichment in several HDAC-regulated gene sets.

### IPMK knockout increases acetylation of histone H4 by proteomic mass spectrometry

Both the enrichment in genes regulated by HDAC pathways in the RNA seq data and the bias towards transcript upregulation in the IKO cells suggested histone acetylation may be increased in IKO cells. We tested this using proteomic mass spectrometry on histones (**Spreadsheet 3**), which showed histone H4 acetylated on lysines 5,8,12 and 16 (H4K5acK8acK12acK16ac) was increased in IKO *vs*. WT cells (**Fig 2A**). To determine any bias toward up-regulation or down-regulation of acetylation, we examined the distribution of changes in all peptides containing any modification (acetylation, methylation and/or phosphorylation), suggesting H4 modifications were significantly biased toward upregulated modifications in IKO relative to WT cells (93% upregulated) compared to histone H3 modified peptides (only 40% upregulated, **Fig 2B**). The H4 bias toward upregulated modifications was also observed when only the acetylated peptides were examined (**Fig 2C**). Using a dot plot to visualize changes in all acetylated peptides with increased modifications (phosphorylated and methylated peptides excluded) shows the largest changes to acetylation occurred in histone H4 peptides (**Fig 2D**), and pair-distances between upregulated acetylated peptides highlights that although most peptides had broadly similar responses to *IPMK*-knockout (**Fig 2E, Fig S2**), the largest differences were in acetylated H4 peptides (**Fig 2D**). These data suggest that *IPMK* loss increases acetylation of histone H4, as determined by proteomic histone mass spectrometry.

**Figure 2.**
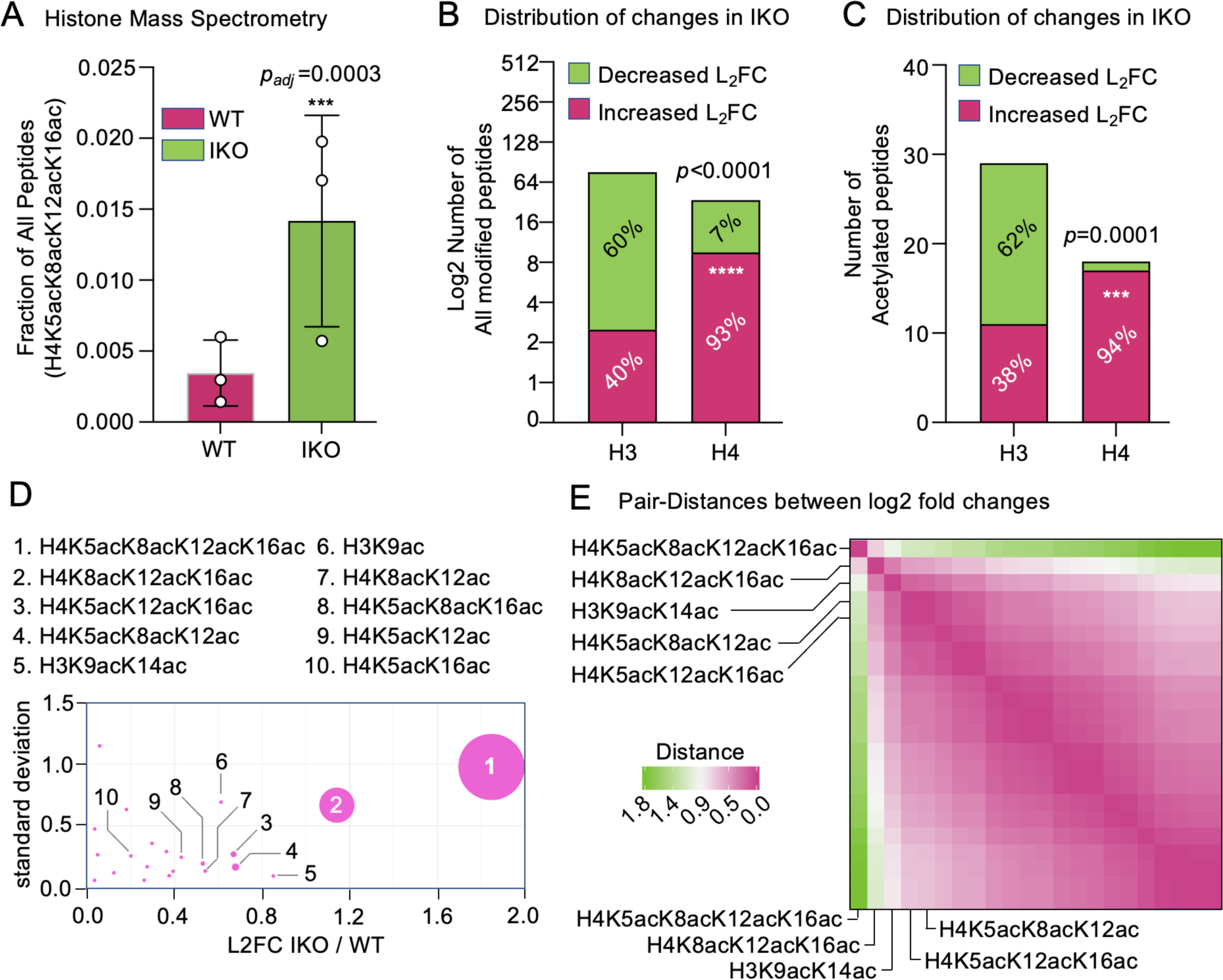
Mass spectrometry of histones shows IPMK knockout upregulates histone H4 acetylation. **A.** Histone proteomic mass spectrometry from triplicate samples of the H4K5acK8acK12acK16ac peptide as a fraction of all detectable peptides in IKO vs. WT U251 cells, two-way ANOVA test with Sidak’s correction, n=3. **B**. Contingency analysis of all modified (acetylated, phosphorylated and/or methylated) histone H3 and H4 peptides, comparing enrichment of peptides with log2 fold changes that increased (green) *vs*. decreased (red) in IKO cells, *p*<0.0001 by Fisher’s exact, these data suggest modifications are more frequently increased on histone H4 *vs.* histone H3. **C**. Contingency analysis of only acetylated histone H3 *vs.* H4 peptides (phosphorylated and methylated peptides excluded), comparing enrichment of peptides with log2 fold changes that increased *vs*. decreased in IKO cells, *p=*0.0001 by Fisher’s exact, suggesting acetylation marks are more frequently increased on histone H4 *vs.* histone H3. **D**. Dot plot of all quantified peptides with acetylation modifications and increased log2 fold changes in IKO cells (phosphorylated and methylated peptides excluded), dot size indicates –log(*p_adj_*) value from two-way ANOVA with Sidak’s correction, peptides of interest are labeled 1 through 10 as indicated. These data illustrate that peptides 1 and 2 are the most changed by IPMK knockout. **E**. Pair-distances of log2 fold changes of all modified histone H3 and H4 peptides comparing WT vs. IKO, suggesting the most different peptides between WT vs. IKO are highly acetylated H4 peptides; complete labels for all peptides on the plot are provided in supplemental data. These data suggest IPMK knockout increases histone H4 acetylation, as measured by proteomic histone mass spectrometry.

### IPMK knockout increases H4-acetylation by ChIP-seq

To further investigate if loss of IPMK may upregulate histone H4 acetylation, we carried out ChIP-seq of poly-acetylated histone H4 (H4ac) (**Spreadsheet 4**). DiffBind identified 31,101 loci with differential H4Ac underlying count data in IKO vs. WT chromatin (FDR<0.05, **Spreadsheet 5**). Decreased acetylation in IKO cells was observed at 10,816 loci (34.8%) but H4Ac was increased at 20,285 loci (65.2%), suggesting IKO increases H4-acetylation (**Fig 3A**), consistent with the proteomic mass spectrometry (**Fig 2**). Compared to wild-type, H4Ac ChIP signal in IKO cells was enriched adjacent to transcriptional start sites, with maxima at −240bp and +190bp relative to the start site (**Fig 3B**, gray regions). H4Ac fold enrichment near the start sites of genes differentially expressed in the IKO transcriptome studies (**Fig 1F**) showed increased acetylation in IKO relative to WT cells (**Fig 3C**). We then binned the data into start sites of transcripts either downregulated or upregulated by IKO in the transcriptome studies (**Fig 1**). While H4Ac fold enrichment increased at start sites of transcripts upregulated by IKO as expected, no difference in H4Ac enrichment was observed at start sites of transcripts downregulated by IKO (**Fig 3D-F**). Together, these data suggest loss of IPMK affects gene expression in two ways, one which increased transcript abundance and H4Ac at those start sites, and another decreases transcript abundance without changes to H4Ac at those start sites.

**Figure 3.**
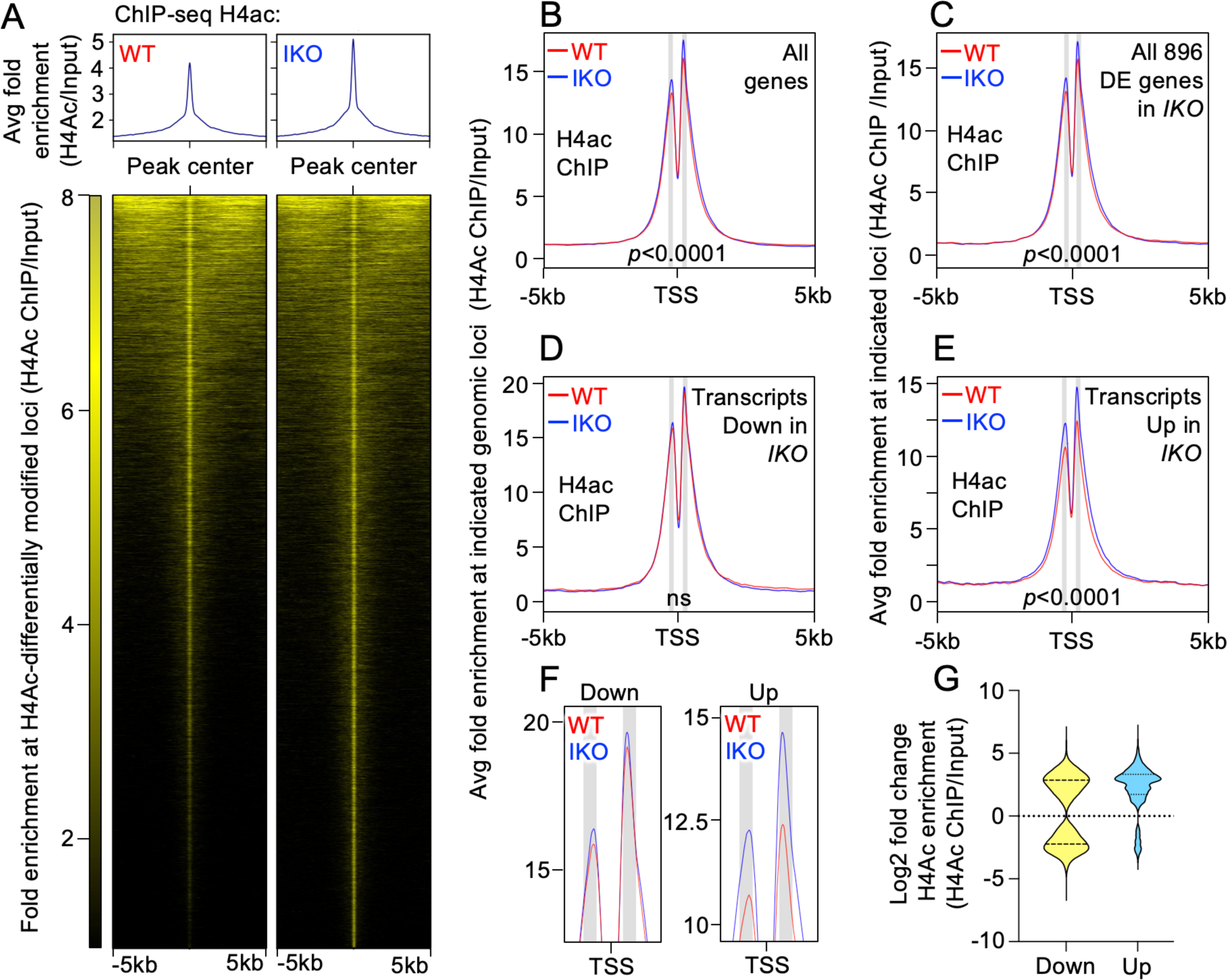
ChIP-seq shows IPMK knockout increases histone H4 acetylation at transcriptional start sites. **A.** Enrichment of anti-poly acetylated histone H4 antibody ChIP-seq signal (H4Ac) in *IKO* cells aligned to peak centers in upper panel A, and heatmap of individual peak sites at all differentially modified (diffbind FDR<0.05) loci in lower panels, suggesting IKO broadly increases H4 acetylation. **B.** Average H4Ac ChIP-seq fold enrichment at all transcription start sites (TSS) comparing WT (red) *vs.* IKO (blue) at all H4Ac differential (diffbind FDR<0.05) refseq genes upon IKO (*p*<0.0001, Fisher exact), greyed regions are peak centers in all panels. **C.** Average H4Ac ChIP-seq fold enrichment at the TSS of all differentially expressed transcripts in IKO *vs*. WT cells (*p_adj_*<0.05), showing increased H4Ac in IKO cells at these sites (*p*<0.0001 Fisher exact). **D.** Average H4Ac ChIP-seq fold enrichment at the TSS of genes whose transcripts were downregulated in IKO *vs*. WT cells (*p_adj_*<0.05), showing no significant (ns) change to H4Ac in IKO cells at these sites. **E.** Average H4Ac ChIP-seq fold enrichment at the TSS of genes whose transcripts were upregulated in IKO *vs*. WT cells (*p_adj_*<0.05), showing increased H4Ac in IKO cells at these sites (*p*<0.0001 Fisher exact). **F.** Enlarged view of peak centers from panels D and E, **G**. violin plots of the log2 fold change of H4Ac ChIP enrichment at the TSS of all transcripts that were either Downregulated by IKO (Down, yellow) or Upregulated by IKO (Up, blue), showing bias towards increased H4Ac signal occurs at the start sites for upregulated transcripts. These data suggest IPMK-knockout increases H4Ac by ChIP-seq at transcriptional start sites.

### HDAC target genes are regulated in an IPMK kinase-dependent manner

IPMK has both kinase-dependent and kinase-independent roles in transcription^20,30^, to clarify the role of IPMK kinase activity in the observed gene expression changes, we complemented IKO cells with wild-type HA-IPMK (IKO*+*WT*-*IPMK), a kinase dead IPMK (D144A) (IKO*+*KD*-*IPMK), or empty vector (IKO*+*Vector) using recombinant retroviruses to generate cell lines expressing these proteins (**Fig 4A**). Wild-type IPMK expression rescued higher-order inositol phosphate production as expected and to similar levels observed in studies from other groups^34^, while kinase-dead IPMK failed to rescue (**Fig 4B**). Wild-type IPMK expression rescued all IKO growth defects, while kinase-dead IPMK cell growth was indistinguishable from IKO (**Fig 4C**). RNA-seq showed that wild-type IPMK expression in IKO cells results in differential regulation of 1,903 transcripts (*p_adj_*<0.05) or 308 transcripts (*p_adj_*<0.05 and Log_2_FC±1.0) in an IPMK-kinase dependent manner (**Fig 4D** and **Spreadsheet 6**) while kinase-dead IPMK only changed abundance of 40 transcripts (*p_adj_*<0.05) or 10 transcripts (*p_adj_*<0.05 and Log_2_FC±1.0) (**Fig 4E**). Gene set enrichment of the IPMK-kinase dependent genes again discovered significant overlap suggest the kinase activity of IPMK controls expression of genes that significantly overlap with HDAC-regulated gene sets, again consistent with IPMK kinase-activity activating HDAC-deacetylase enzyme activity.

**Figure 4.**
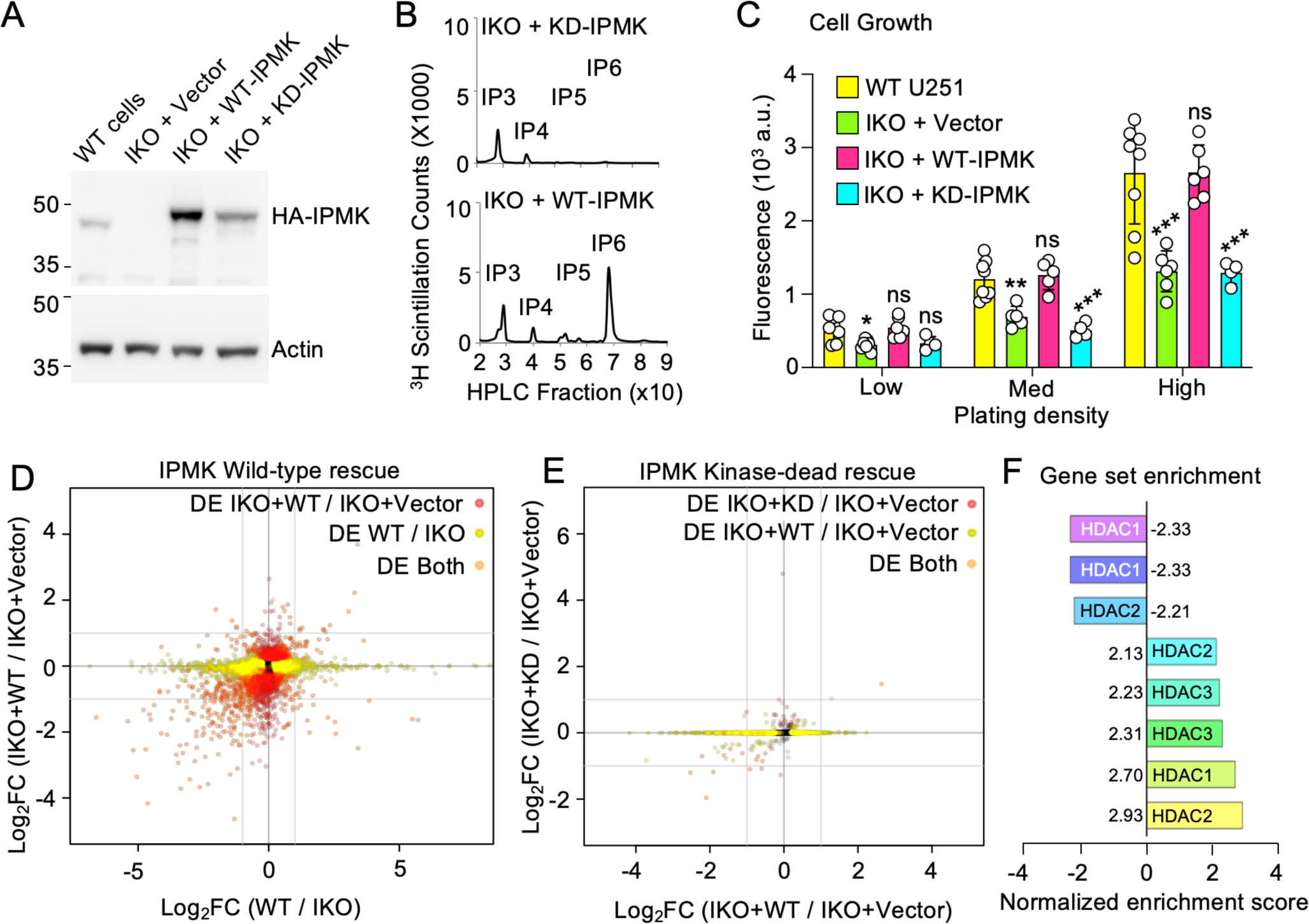
The IPMK kinase-dependent transcriptome is enriched in HDAC-regulated genes. **A.** Representative western blot of indicated transduced cell lines, slower migration of expressed IPMK band likely due to the N-terminal HA-tag. **B.** HPLC of metabolically ^3^H-inositol radiolabeled phospho-inositols from indicated U251 cell lines, showing peaks identified using standards for IP3, IP4, IP5 and IP6. As expected, WT-IPMK increased IP5 and IP6 levels while KD-IPMK had no effect. **C.** CyQuant DNA content cell growth assay of indicated cell lines, plated at indicated cell densities with DNA content measured at 72 hours, \**p_adj_*<0.05, \*\**p_adj_*<0.01, \*\*\**p_adj_*<0.001 in ordinary one-way ANOVA Dunnett’s corrected; ns is not significant. **D.** Volcano plot of log2 fold changes of differentially expressed (*p_adj_*<0.05) transcripts in WT *vs.* IKO cells (y-axis) and WT-IPMK *vs.* vector rescue in IPMK-knockout (IKO) cells (x-axis), showing 308 genes (*p_adj_*<0.05 and Log_2_FC±1.0, orange dots) or 1903 genes (*p_adj_*<0.05 only, yellow dots) differentially expressed. **E.** Same as in D, but x-axis is kinase-dead KD-IPMK *vs*. vector rescue in IKO cells, showing only 10 genes (*p_adj_*<0.05 and Log_2_FC±1.0, orange dots) or 40 genes (*p_adj_*<0.05 only, yellow dots) as differently expressed. **F.** Normalized gene set enrichment scores of HDAC gene sets from the IPMK kinase-dependent transcriptome (1903 genes, *p_adj_*<0.05), full results provided as a supplemental spreadsheet, GSEA used the Molecular Signatures database. These data suggest the kinase activity of IPMK regulates gene expression in U251 cells, and those differently expressed genes are significantly enriched in HDAC-regulated genes.

### IPMK kinase activity or exogenous IP4 is required for full deacetylase activity of HDAC3 immuno-precipitates

Inositol phosphates have been co-crystalized with HDAC1^36^ and HDAC3^37^ (**Fig 5A**) and many diverse inositol phosphates have been shown to activate purified Class 1 HDACs tested *in vitro*^6,7^. We therefore tested if HDAC enzyme activity in human cells is dependent on cellular inositol phosphates in IKO cells. We observed no significant changes to Class 1 HDAC transcripts in IKO cells (**Fig 5B**), consistent with IPMK controlling Class 1 HDAC enzyme activity, but not expression levels^38^. We then immunoprecipitated endogenous, untagged HDAC enzymes from WT or IKO native whole cell extracts using isoform-selective antibodies, and monitored HDAC deacetylase enzyme activity by *in vitro* HDAC deacetylase assays. No changes between WT and IKO cells were observed in HDAC activity present in immunoprecipitates using HDAC1, HDAC2 or HDAC8 antibodies, but immunoprecipitates of HDAC3 showed significantly decreased HDAC activity from IKO cells (**Fig 5C-F**). The signal in this enzyme assay was reduced to background levels in the presence of Trichostatin A (**Fig S3**). No changes in HDAC3 protein levels were observed in inputs or the HDAC3 immunoprecipitates by westerns (**Fig 5G, Fig S6-S7**) and no changes were observed in HDAC3 transcript abundance (**Fig 5B**). HDAC3 activity immunoprecipitated from IKO cells expressing recombinant wild-type IPMK was fully rescued, while kinase-dead IPMK expression had no effect on enzyme activity in HDAC3 immunoprecipitates (**Fig 5H**), suggesting IPMK kinase activity is required for full HDAC3 deacetylase activity. The HDAC3 enzyme was previously co-crystalized with endogenous Ins(1,4,5,6)P4 that co-purified with HDAC3 from animal cells^11^ (**Fig 5A**), suggesting this species associates with native HDAC3. Abundance of this IP4 species Ins(1,4,5,6)P4 has been shown to depend on the kinase activity of IPMK^39–42^. Addition of chemically synthesized Ins(1,4,5,6)P4 to the *in vitro* deacetylase reaction fully rescued the defect in HDAC3 immunoprecipitates from IKO cells (**Fig 5I**), while control inositol had no effect on the HDAC3 enzyme reactions (**Fig 5J**). The addition of Ins(1,4,5,6)P4 did not detectably change the pH of the HDAC enzyme reaction (**Fig S5**). These data suggest the kinase activity of IPMK is required for full deacetylase enzyme activity of immunopurified HDAC3, and this defect can be fully rescued by addition of chemically synthesized Ins(1,4,5,6)P4 to the HDAC reaction.

**Figure 5.**
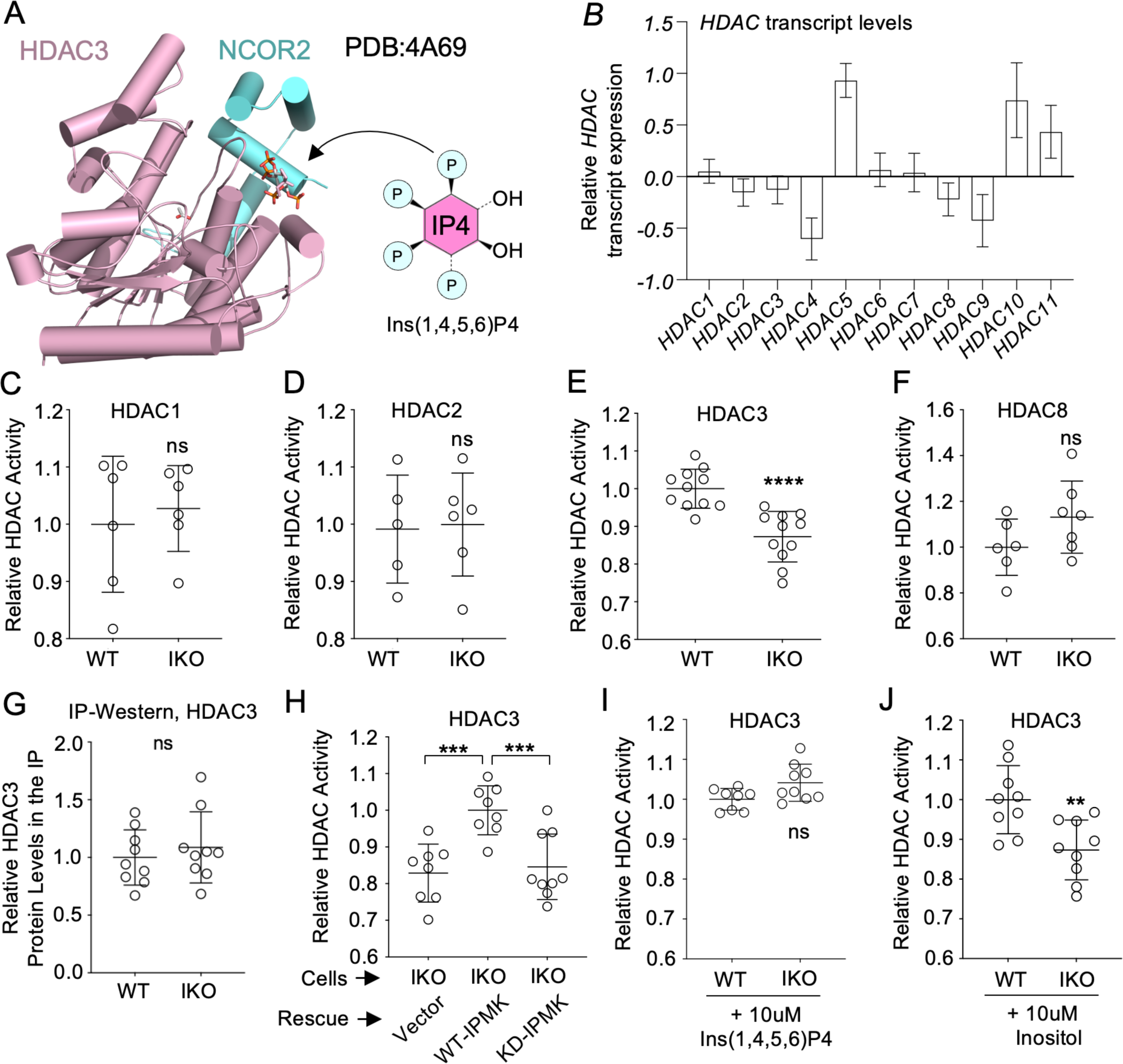
Full HDAC3 enzyme activity depends on the kinase activity of IPMK. **A.** Crystal structure (PDB:4A69) of HDAC3 co-crystalized with Ins(1,4,5,6)P4. **B.** RNA-seq mRNA transcript levels of all HDACs in IPMK-knockout (IKO) cells relative to WT cells. **C,D,E,F.** Relative HDAC deacetylase enzyme activities contained in HDAC1, HDAC2, HDAC3 and HDAC8 immunoprecipitates from native whole cell extracts, \*\*\*\**p*<0.0001 by unpaired t-test, ns is not significant. **G.** Quantitation of triplicate anti-HDAC3 Westerns of anti-HDAC3 immunoprecipitates from native whole cell extracts, showing no decrease in HDAC3 protein levels by t-test (ns). Total HDAC3 in the inputs was also unchanged by Western (see supplemental data for all full westerns and exposures) **H.** HDAC deacetylase enzyme activity contained in HDAC3 immunoprecipitates from native whole cell extracts of *IKO* cells expressing either vector, wild type (WT-IPMK) or kinase-dead (KD-IPMK), showing WT-IPMK rescues deacetylase activity but kinase-dead KD-IPMK does not, *\*\*\*p<*0.001 by unpaired t-test **I.** Relative HDAC deacetylase enzyme activity contained in HDAC3 immunoprecipitates from WT or IKO extracts in the presence of chemically synthesized Ins(1,4,5,6)P4 (10uM) depicted in panel A above or **J.** control inositol (10uM) added back to the HDAC enzyme assay, these data suggest Ins(1,4,5,6)P4 restores HDAC3 activity, pH was confirmed identical before and after addition of inositol phosphates to these enzyme reactions (see supplemental data). These data suggest HDAC enzyme activity is decreased in HDAC3 immunoprecipitates from IPMK-knockout cells, which can be rescued either by expression of wild-type IPMK but not kinase-dead IPMK, or by addition of chemically synthesized Ins(1,4,5,6)P4 to the assay.

## DISCUSSION

Data presented here support a model in which the kinase activity of IPMK is required for production of certain inositol phosphates (**Fig 6A**) which activate HDAC3 in human cells (**Fig 6B**). We did not observe changes to Class 1 HDAC protein levels or transcript levels upon IPMK knockout, consistent with results from the brain-specific knockout of mouse *Ipmk*, in which Class 1 HDAC protein levels also did not change^38^. Here, *IPMK* knockout decreased HDAC3 enzyme activity and increased histone H4 acetylation. ChIP-seq data suggests histone H4 acetylation increased at genes upregulated by IPMK loss, while genes downregulated by IPMK loss had no change in H4 acetylation. A limitation of this study is that we cannot know the fraction of the increased H4 acetylation that can be assigned exclusively to the decrease in HDAC3 activity (**Fig 6C-D**). It is likely that some fraction of the increased H4 acetylation is independent of HDAC3, as IPMK is known to regulate a particularly wide array of cellular pathways established by other studies^33,38,43–47^. Nevertheless, the data here directly support genetic studies in yeast which demonstrate deletion of the yeast HDAC orthologue (Rpd3L) causes disruption of stress-induced transcriptional changes that were phenocopied by loss of inositol phosphate synthesis enzymes^48^. Thus, evidence from multiple labs and multiple model systems all suggest the increase in H4 acetylation observed here is in part due to a decrease in HDAC3 activity.

**Figure 6.**
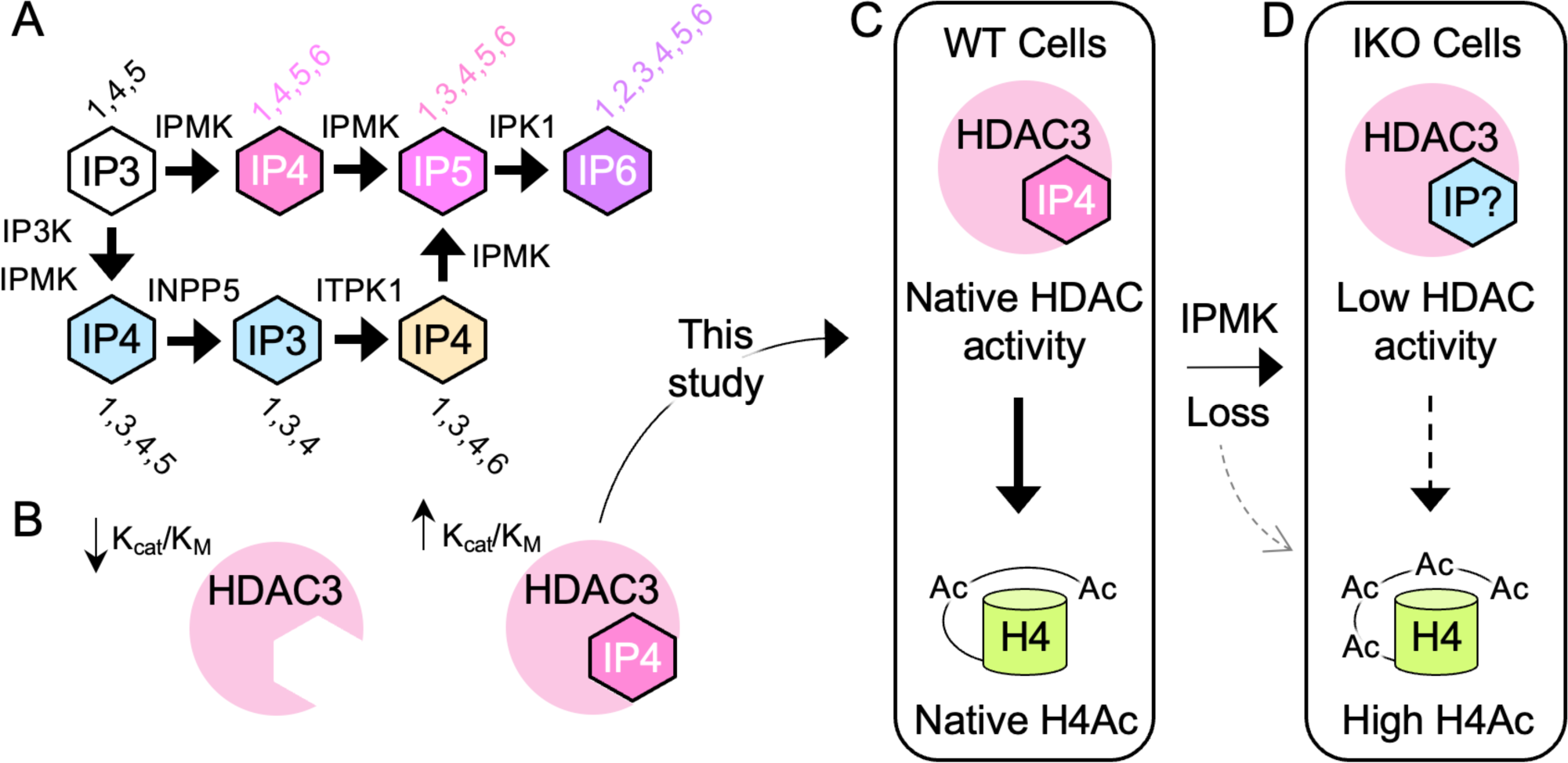
Model of IPMK-controlled HDAC3 activity. **A.** This study and previous studies have shown IPMK loss removes all detectable IP5 and IP6. However, some IP4 species, namely Ins(1,3,4,5)P4 (blue IP4) and Ins(1,3,4,6)P4 (yellow IP4) can be synthesized by other enzymes and are expected to persist in IPMK knockout cells, consistent with studies here (Fig 1). **B**. HDAC3 can co-purify and co-crystalize with Ins(1,4,5,6)P4, which activates HDAC3 enzyme activity, while Ins(1,3,4,6)P4 was shown in other studies to not activate HDAC1 or HDAC3 enzyme activities *in vitro*^7^. **C.** In this study, we deleted IPMK from human U251 glioblastoma cells, which altered native HDAC3 enzyme activity and histone H4-acetylation. **D.** A limitation of this study is that the precise fraction of the observed increase in H4-acetylation that is due directly to loss of HDAC3 activity is unclear, as genetic loss of IPMK could affect other mechanisms that regulate histone acetylation in living cells (gray dashed arrow). We conclude that IPMK loss leads to 1) decreased HDAC3 activity and 2) increased H4-acetylation, in human U251 glioblastoma cells.

Perhaps most intriguing is why IPMK loss did not affect the activity of all Class I HDACs, since extensive previous studies have shown HDAC1 and HDAC3 both have increased k_cat_/K_M_ in the presence of IP5 and IP6^49^, and both IP5 and IP6 are decreased to undetectable levels in IPMK-knockout cells (**Fig 1**). However, levels of total IP4 do not change upon IPMK knockout, as has been observed in other studies^34^. The remaining IP4 likely represents two IP4 species that can be produced through IPMK-independent pathways (**Fig 6A**). Previous studies have demonstrated Ins(1,3,4,6)P4 does not activate HDAC1 or HDAC3, but Ins(1,3,4,5)P4 potently activates both HDAC1 and HDAC3^49^. Thus, the simplest hypothesis is that the persistent IP4 species in IKO cells is sufficient to maintain HDAC1 activity, but is insufficient to maintain HDAC3 activity. This is further supported by extensive enzyme kinetic, direct binding and computational docking studies^49^ that suggest Ins(1,3,4,5)P4 does not activate HDAC3 as robustly as Ins(1,4,5,6)P4 due to the stereochemistry of the inositol ring, although multiple inositol phosphates effectively activate HDAC3, such as Ins(1,4,5,6)P4 (k_cat_/K_M_=0.0326 s^-1^μM^-1^) and IP6 (k_cat_/K_M_=0.0346 s^-1^μM^-1^)^49^. Nevertheless, an inherent limitation of this study, and any study that attempts a comparison of cell-based genetics to enzyme kinetics, is that any number of cellular events could be responsible for the decreased HDAC3 activity. Indeed, a recent study in mouse intestinal epithelial cells suggested extracellular Ins(1,4,5)P3 can activate intracellular HDAC3^14^, despite a great deal of *in vitro* data suggesting Ins(1,4,5)P3 can neither bind nor activate HDAC3^49^, highlighting the potential complexity of indirect effects in living cells. Here, we demonstrate ectopic, chemically synthesized Ins(1,4,5,6)P4 completely rescued deacetylase activity in HDAC3 immunoprecipitates from IPMK-knockout cells, suggesting Ins(1,4,5,6)P4 as a potential cellular factor required for full HDAC3 deacetylase activity. The particular inositol phosphate species required for full activity of HDACs has not been established, although the complexity of the signaling suggests even genetic analyses of other inositol phosphate signaling enzymes in parallel (including *IP3K*, *INPP5*, *ITPK1* and *IPK1*) may be difficult to interpret. Chemical inhibitors of IPMK could be used to interrogate the detailed requirements of HDAC1, HDAC2 or HDAC8 for specific inositol phosphate species, selective and potent inhibitors of IPMK are currently under development^50–52^ could aid in further interrogating the specificity of IPMK regulation of HDAC3.

Regardless of how IPMK loss is regulating HDAC3, that HDAC3 was selectively regulated could have therapeutic implications for several human pathologies. Several studies have reported specific roles for HDAC3 in solid tumors^53^, rare childhood cancers^54^, prostate cancers^55^, spinal cord injuries^56^ and in diet-induced obesity^57^. All these pathologies could potentially benefit from specific inhibition of HDAC3. IPMK is a kinase with a typical ATP-binding site, several groups are currently developing IPMK inhibitors^50–52^. The connection to HDAC3 activity in human cells imparts new potential value to these IPMK kinase inhibitors, with potential to help increase the specificity of HDAC3-targeted interventions. In addition to those more clinical implications, the work here brings new basic science questions to light regarding how IPMK regulates gene expression. Since H4-acetylation was increased at genes upregulated by IKO, but unchanged at genes downregulated by IKO, the data are consistent with IPMK using a second mechanism to regulate gene expression that is not currently described, and appears to be independent of H4 acetylation. More studies will be needed to elucidate those details, however we previously published a mechanism wherein IPMK directly phosphorylated a phosphoinositide lipid bound to a transcription factor, in which loss of IPMK decreased gene expression mediated by that transcription factor^33^, similar to the phenotype of IPMK loss observed here. Further study will be required to investigate the mechanisms used by IPMK to control gene expression, and the implications in human disease.

## ACKNOWLEGMENTS

Support was from the National Institutes of General Medical Sciences (NIGMS) R01 GM132592 to RDB. We thank members of the Blind Lab and Prof. John D. York for helpful discussions, Prof. Ashok Venkitaraman for IPMK antibody, EAS was supported by T32 DK007563, JCP was supported by T32 DK101003 - DK020593; ATH was supported by F30 HL143826. We acknowledge assistance from the Center for Innovative Technology at Vanderbilt University for Mass Spectrometry; The Vanderbilt Mass Spectrometry Resource Center, Vanderbilt Ingram Cancer Center P30 CA68485 and The Vanderbilt Institute of Chemical Biology.

## MATERIALS and METHODS

### Materials

Human glioblastoma U251-MG cells were obtained from ECACC #09063001, IPMK antibody was a gift from Prof. Ashok Venkitaraman^58^, HDAC1 antibody Cell Signaling Technology #34589; HDAC2 antibody Cell Signaling Technology #57156, HDAC3 antibody Cell Signaling Technology #85057, HDAC8 antibody Cell Signaling Technology #85057, H4ac Pan-acetylation antibody Active Motif #39243, Protein-G magnetic beads Cell Signaling Technology #9006, HDAC deacetylase assay Sigma #CS1010, D-*myo*-inositol-1,4,5,6-tetraphosphate (Ins(1,4,5,6)P4) Cayman Chemical #10007783, and D-*myo*-inositol MP Biomedicals #102052.

### IPMK genetic deletion

CRISPR-Cas9-mediated genetic deletion of all copies of human *IPMK* in human U251-MG cells (ECACC 09063001) was performed as previously described^59,60^. Briefly, guide RNAs were optimized and selected by the computational tool “CRISPOR”^61^ to enhance target selectivity for *IPMK* exons 1 and 6 (see Supplemental for gRNA locations at the *IPMK* locus). IPMK guide RNA sequences for Exon 1 were 5’-CACCGGCGATCGAGTCCACCCCTGA-3’ (A, Forward) and 5’-AAACTCAGGGGTGGACTCGATCGCC-3’ (B, Reverse). Exon 6 guide RNAs were 5’-CACCGCCAAGATGTATGCGCGTCAC-3” (C, Forward) and 5’-AAAC GTGACGCGCATACATCTTGGC-3’ (D, Reverse). Oligonucleotide pairs “A/B” and “C/D” were homologous to portions of *IPMK* exon 1 and 6, respectively, these pairs were annealed and cloned into the BbsI site of pX459v2^62^, resulting pX459v2-*IPMK*-exon1 and pX459v2-*IPMK*-exon6, which were co-transfected into human U251-MG glioblastoma cells (ECACC 09063001) using Lipofectamine 3000 (ThermoFisher). Transfectants were selected for three days at which point selection antibiotic was removed, followed by single cell cloning in 96 well plates. Wells containing one colony were expanded and screened for IPMK knockout (IKO) by western blot. IKO was confirmed by analytical PCR using genomic extracted DNA as shown in supplemental, the sequence of all alleles was determined by Sanger sequencing^60^.

### Inositol phosphate ^3^H-metabolic labeling and HPLC

All cell lines (WT, IKO, and IKO expressing wild type N-terminally HA-tagged IPMK, or the kinase-dead HA-IPMK mutant D144A^63^) were grown in DMEM supplemented with 10% FBS, non-essential amino acids and penicillin/streptomycin, medium was changed to inositol-free media (RPMI) and cells were inositol-starved for 1 day. Cells were maintained in culture containing inositol-free DMEM medium and supplemented with 100µCi ^3^H-inositol (Perkin Elmer) for 5 days. Cells were then washed in PBS, manually scraped from the dish and collected in 100µL HCl. Inositol phosphates (IPs) were extracted as described previously^64^, by adding 372µL of 1:2 CHCl3:MeOH to one well of a 6-well plate (approximately 10^6^ cells), vortexed and followed by 125µL CHCl3 and 125µL 2M KCl added to each sample, vortexed and centrifuged 5 minutes at 14,000 rpm to separate upper aqueous and lower organic phases, upper phase contains soluble IPs. HPLC analysis of the upper phase used a Partisphere SAX 5 4.6 x 125 mm column, eluted with 10mM (Buffer A) to 1.7M (buffer B) ammonium phosphate (pH= 3.5). 1mL fractions were collected and mixed with 6mL scintillation fluid and DPM measured by a liquid scintillation counter. Elution of IP3, IP4, IP5 and IP6 standards confirmed the identity of each peak off the HPLC, IP data is normalized to total DPM counts, quantification represents three independent metabolic labeling experiments with error representing standard error of the mean.

### Expression of wild type or kinase-dead IPMK

N-terminally HA-tagged IPMK (WT or the kinase deficient D144A mutant^63^) were cloned into pLPCX (Clontech). VSVg pseudotyped MLV was prepared by transfecting pLPCX-HA-IPMK along with pMLV-Gag-Pol and pVSVg into HEK293T cells using PolyJet transfection reagent (SignaGen). Released virus was harvested and HA-IPMK WT or D144A viruses were used to infect IPMK-knockout (IKO) human U251-MG cells (clone 2-F1) at MOI of less than 1 as previously described^59^. Control cells were transduced with empty vector, selection for puromycin-resistance was initiated at 3 days post infection after which cells were maintained in puromycin containing medium. Cells were tested by western blot for expression of the IPMK in whole cell extracts.

### RNA-seq and gene set enrichment analysis

DNase-treated RNA prepped from one well of a six well plate (approximately 10^5^ cells) was extracted using Quick-RNA MiniPrep (Zymo Research) from indicated cells. Cells used for RNA-seq were 1) wild-type U251-MG cells (ECACC 09063001), 2) monoclonal *IPMK*-knockout U251-MG cells (IKO), and IKO cells that were transduced with 3) empty vector-, 4) wild-type HA-IPMK-, or 5) kinase-dead HA-IPMK (D144A)-expressing recombinant MLV retroviruses. PolyA selected RNA-seq libraries were prepared and sequenced at the Vanderbilt Technologies for Advanced Genomics (VANTAGE) core on an Illumina NextSeq500, bases and reads with low quality scores or reads, respectively, were removed and Illumina library adapter sequences were trimmed using bbDuk (https://jgi.doe.gov/data-and-tools/software-tools/bbtools/bb-tools-user-guide/bbduk-guide/). The resulting files were mapped to hg38 using HiSat2^65^, poorly mapped reads were removed with SamTools^66^. Reads mapped to RefSeq genes were counted using HTSeq-count^67^ and differential expression assessed using DESeq2^68^ with Apeglm LFCShrinkage^69^. Gene sets from the molecular signatures database (version 6.1) were used to apply SetRank ^70^ and Camera^71^ provided as supplemental spreadsheets, all original FASTQ files and underlying count data are available upon request.

### Histone mass spectrometry

Histones from the nuclei of indicated U251-MG cells were acid extracted and histones H3 and H4 were SDS-PAGE gel purified and subjected to LC/MS/MS as previously described, 4N H_2_SO_4_ was added to cell nuclei pellets and incubated at 4°C, the extracts centrifuged, and supernatant histones extracted again to pellet traces. TCA (20% final) was added to the histones, incubated on ice and centrifuged. Histone pellet/film was washed with acetone + 0.1% HCl, centrifuged for 5 min, the histone pellet washed twice with 100% acetone and allowed to dry. The histone film was resuspended in water and centrifuged to pellet any remaining debris. Histones were run on SDS-PAGE to separate histone H3 and H4 bands. Coomassie-stained bands containing Histone H4 were excised, diced into 1mm^3^ cubes, and gel pieces destained in 50% acetonitrile in 25mM ammonium bicarbonate. Destaining solution was removed, gel pieces were washed twice with 100mM ammonium bicarbonate, and excess buffer was removed. Histone H4 was chemically derivatized and digested using similar methods as previously described^72^. Derivatization was accomplished by addition of propionylation reagent (propionic anhydride and anhydrous methanol in a ratio of 3:1). Gel pieces were incubated for 20 minutes at room temperature, propionylation reagent was removed, gel pieces were rinsed with 100mM ammonium bicarbonate, and derivatization was repeated. After derivatization, 75% acetonitrile was added to gel pieces, incubated for 5 minutes, and acetonitrile removed. Proteins were then in-gel digested with trypsin (10ng/μL) in 25mM ammonium bicarbonate overnight at 37°C. Peptides were extracted from the gel by addition of 60% acetonitrile and dried by speed vac centrifugation. Peptides were reconstituted in 20μL of 100mM ammonium bicarbonate and derivatized by two consecutive reactions with propionylation reagent in order to fully derivatize the amino-termini of histone peptides. Propionylation reagent (10μL) was added to reconstituted histone peptides, followed by addition of concentrated ammonium hydroxide to re-establish pH 8 for derivatization. Derivatization was performed for 15 minutes at 37°C, samples were dried, and peptides were reconstituted in 0.1% formic acid for analysis by LC-coupled tandem mass spectrometry (LC-MS/MS). Peptides were loaded onto a reverse phase analytical column, 360μm O.D. x 100 μm I.D. fused silica packed with 3μm Jupiter C18 reverse phase material (Phenomenex), using a Dionex Ultimate 3000 nanoLC and autosampler. Mobile phases consisted of 0.1% formic acid, 99.9% water (solvent A) and 0.1% formic acid, 99.9% acetonitrile (solvent B). Peptides were gradient-eluted at a flow rate of 350 nL/min using a 75-minute gradient. The gradient consisted of 0-25%B in 58 minutes; 25-90%B in 4 minutes; 90%B for 1 minute; 90-0%B in 2 minutes; and 0%B for 10 minutes (column re-equilibration). Peptides were analyzed on a Q Exactive Plus mass spectrometer (Thermo Scientific), equipped with a nanoelectrospray ionization source. The data acquisition method consisted of both data-dependent and targeted scan events. Full scan spectra (m/z 300-1400) were acquired with an AGC target of 3e6, followed by 12 DDA MS/MS scans of the most abundant ions detected in the preceding MS scan, and 3 targeted MS/MS scan events using an AGC target of 5e4. Targeted MS/MS were acquired for specific m/z values (m/z 768.9465, 761.9386, 754.93) corresponding to isobaric species of acetylated H4 peptide 4-17. HCD collision energy was set to 27 nce. Histone H4 post-translation modifications were identified and quantified using EpiProfile (2.0) which first identified PTMs by retention time relationships between modified and unmodified peptides (layouts) and then quantified the PTMs by determining the area under the curve (ratios). Ratios were used to calculate log2 fold changes and statistical differences between modified peptides from WT and IKO histone by two-way ANOVA test with Sidak’s correction for multiple comparisons.

### Chromatin Immunoprecipitation

For each ChIP experiment duplicate 15cm plates containing at least 2.5×10^6^ of the indicated U251-MG derived cell lines were cross linked with 1% formaldehyde from a freshly opened sealed glass ampule for 10 minutes at room temperature, quenched with 2.5 M glycine (125 mM final), the cells washed 3 times with PBS to remove media^73^. Following aspiration of PBS, 5 mL of Farnham lysis buffer (5 mM PIPES pH 8.0, 85 mM KCl, 0.5% NP-40, EDTA-free Roche protease inhibitor tablets, 10mM Sodium butyrate (pH 7.0)) was added per plate. Cells were scraped, pelleted, and flash frozen in liquid nitrogen. Frozen cell pellets were resuspended 1mL of Farnham lysis buffer per 2.5×10^6^ cells, rocked 15 min at 4°C, then dounced 50 times with a type B pestle. The crude preparation of nuclei was pelleted at 600xg for 5 minutes at 4°C, the pellet resuspended in 300 µL of Nuclear Lysis Buffer per 2.5×10^6^ cells (50 mM Tris pH 8.0, 10 mM EDTA, 0.5% SDS, and 1% NP40, 10mM Sodium butyrate (pH 7.0)) and placed in a 1.5mL Diagenode polystyrene sonication tubes. Chromatin was sheared in a Bioruptor Pico for ten cycles of 30 sec on, 30 sec off until fixed lysate is clear. Sonicated samples were spun at 16,000XG for 15minutes at 4C, 300uL of supernatant was diluted to 1.4mL with ChIP dilution buffer (10mM Sodium butyrate (pH 7.0), 50mM Tris (pH7.5), 1mM EDTA, 1% NP-40, 150mM NaCl, 0.25% Sodium Deoxycholate, protease inhibitors), and snap frozen at −80C. Fragment size was verified by agarose gel electrophoresis to be about 200bp prior to ChIP. ChIP was carried out overnight at 4°C using IgG control or H4ac Pan-acetylation antibody (Active Motif # 39243) with anti-rabbit Dynabeads blocked with BSA. The following day, the beads were washed 2x in low salt (20mM Tris (pH 7.5) 150mM NaCl 0.1% Triton X100, 0.1% SDS, 1mM EDTA) and 2x in high salt wash (20mM Tris (pH 7.5) 500mM NaCl 0.1% Triton X100, 0.1% SDS, 1mM EDTA), 1x with High salt STR buffer (20mM Tris (pH 7.5) 500mM NaCl 1.0% Triton X100, 0.1% SDS, 1mM EDTA), and lastly with Tris/EDTA (TE) buffer. The pellet was resuspended in IP elution buffer (1% SDS, 0.1M NaHCO3), treated with 300ug/mL Proteinase K at a 55°C for 1 hour, crosslinks reversed at 65°C overnight. 150ug/mL RNAseA was added for 1 hour at 37C and total chromatin immunoprecipitated DNA isolated using a Qiagen PCR purification kit and subjected to library prep and sequencing.

### ChIP-seq analysis

Duplicate H4-panAc ChIP DNA and inputs were sequenced by the Vanderbilt VANTAGE sequencing core, libraries were sequenced on an Illumina NextSeq 500 with a minimum of 15 million reads per sample, lowest reads for any sample was 15,617,190 reads for one of the H4panAC ChIP 2-F1*-*IKO samples, all other samples had at least 23 million reads and the average read depth was 27.3 million reads. QC of raw reads was tested using FastQC (0.11.9), low quality reads were removed and Illumina adapter sequences trimmed using bbDuk. Trimmed sequences were then mapped to hg38 using HiSat2 and a sam file generated, poorly mapped and unaligned sequences removed using Hisat2 and SamTools^74^. Sam files containing aligned reads were converted to bam files and duplicate alignments removed using samtools. The unique bam file with duplicates removed was sorted and macs2 used to call the peaks. Peaks were visualized in IgV genome browser. The macs2 bdgcmp utility was used in the pileup.bdg output file from macs2 to reduce noise. Peaks from macs2 were annotated using Homer (annotatePeaks.pl) and the ChIPseeker package in R using bed files to visualize peak coverage over chromosomes and peak profiles at transcriptional start site (TSS) regions. Homer was also used for motif analysis. Differences in H4-panAc ChIP-seq signals between IKO relative to WT were determined by the DiffBind package in R to identify loci with different amounts of underlying counts using the bam file from WT, IKO, inputs and the macs2 narrowPeak files for both WT and IKO conditions. Changes to H4-panAc at the start sites of transcripts differentially expressed by IKO in RNA-seq studies were determined using DiffBind. Briefly, subsets of all quantifiable genes, all IKO differentially expressed genes, the IKO upregulated genes, and *IKO* downregulated genes were made using RefSeq, annotated DiffBind output file was used to obtain fold enrichment scores and total number of differentially bound peaks for each condition above. The differentially bound peaks within each chromatin region as identified by DiffBind were counted and used in contingency analysis to determine statistical enrichment by Fisher’s exact test. Figures were made using bigwig and bed files from WT and IKO compared by compute matrix in Deeptools using the reference point mode. This matrix was used to align peaks by closest transcriptional start site or the peak center and to generate heatmaps and the profile plots. Plot profile was used to create profile plots using the scores from WT and IKO sets over genomic regions in bed files. All FASTQ files are available upon request.

### Cellular proliferation assays

For manual cell counting, equal numbers of cells of the indicated cell lines were plated in triplicate at the indicated cell numbers for each time point tested and were then trypsinized, resuspended in 500uL trypan blue and counted on a hemocytometer using a hand-held manual counter, triplicate hemocytometer counts were averaged. For CyQuant NF cell counting (Thermo Fisher), indicated cell lines were plated equally in a 96-well plate, incubated at 37C, 5% CO2 for 72 hours, each well gently washed with 1X PBS, and CyQuant NF reagent added, incubated for 10min at 37C and 5% CO2, and DNA content in each well measured using a Biotek Neo multimodal plate 384-plate reader at 480nm excitation and 535nm emission, with gain set at 150.

### HDAC immunoprecipitations

Care was taken to maintain native HDAC enzyme activity, no denaturing detergents such as SDS can be used in the immunoprecipitations, any freeze/thaw of HDAC IPs decreased deacetylase activity so immunoprecipitations were prepared and assayed fresh. Indicated cells were grown in media (DMEM with L-glutamine, 4.5g/L glucose, 110mg/L sodium pyruvate, nonessential amino acids, antimycotic and 10% FBS), 5% CO2, 37°C. Since IKO cells had a growth defect, we serum-starved equally seeded IKO and WT cells and reactivated for 6 hours prior to immunoprecipitation to maintain identical growth density. Four million WT or IKO cells were seeded into triplicate 15cm dishes to achieve 70% confluency, the following day the media was replaced with FBS-free media (serum free). After 24 hours cells of this FBS-starvation, the cells were refed with media containing 10% serum for 6 hours, plates washed twice with 1xPBS and scraped into 500ul hypotonic lysis buffer (10mM HEPES pH 7.5, 10mM KCl, 1% triton, EDTA-free protease inhibitors and 10mM beta-mercaptoethanol freshly added), sonicated 1 minute (3 sec on/off) 20% power on a Branson digital probe sonicator with a 1/8inch titanium microtip. NaCl was then added to final concentration of 500mM, and tubes were rocked on ice for 1hr to extract nuclear proteins, then spun at 13,000xg for 30 minutes to produce a cleared supernatant of whole cell extract. Following the high speed spin, high salt from the nuclear extraction was diluted to 166mM with hypotonic lysis buffer, and 1mL of diluted lysate was used for each immunoprecipitation with 5uL of each indicated HDAC antibody (HDAC1 antibody, Cell Signaling Product #34589; HDAC2 antibody Cell Signaling Product #57156, HDAC3 antibody Cell Signaling Product #85057, HDAC8 antibody Cell Signaling Product #85057), with 20uL Protein-G magnetic beads (Cell Signaling Product #9006), rocked overnight at 4C. The following day beads were washed 3x with 700uL of hypotonic lysis buffer and resuspended in 80uL hypotonic lysis buffer + 10mM BME to be used in downstream HDAC enzyme assays. All immunoprecipitates were confirmed to contain equal amounts of each of the indicated HDACs by western blot.

### HDAC deacetylase enzyme assays

Similar to enzyme kinetic studies of Class 1 HDACs^6^, a general HDAC fluorometric deacetylase enzyme assay was used to determine total HDAC activities relative to wild-type U251-MG cells from various HDAC-immunoprecipitates (Sigma CS1010). To each well of a 384-well plate, 4uL of assay buffer and 6uL of each gently resuspended HDAC immunoprecipitation (as described above) was added, the reaction started by addition of 10uL of 20mM fluorometric HDAC substrate and 2uL developer and allowed to proceed for 25 minutes then read on a Biotek Neo multimodal plate 384-plate reader at 360nm excitation and 460nm emission, with the gain set at 150. All raw fluorescence data were normalized by the averaged 460nm emission signal obtained from HDAC IPs from wild-type cells. For IP4 and inositol rescue of HDAC3 enzyme activity, Ins(1,4,5,6)P4 (Cayman Chemical #10007783) or inositol control was added to a final concentration of 10uM, to HDAC3 immunoprecipitated from either wild-type or IPMK-knockout cells, and allowed to bind at 37C for 5 minutes. The HDAC assay was then carried out as described above, we confirmed pH of the complete HDAC reaction was not changed by addition of the IP4 to the HDAC reactions, those data are included in supplemental.

## Supplementary Figure S1

**Figure S1.**
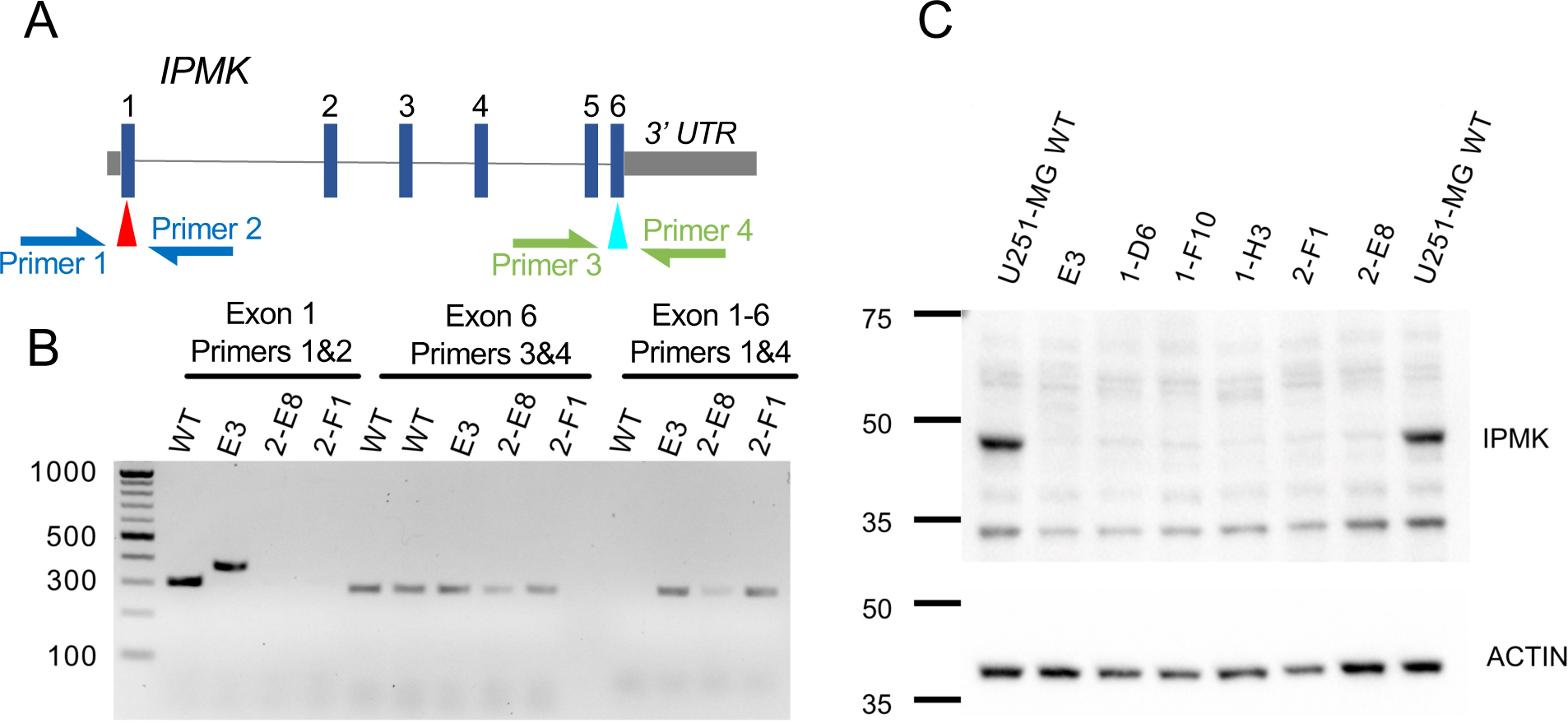
Creation of IPMK knockout cells with CRISPR. **A.** Model of human *IPMK* genetic locus, showing the intron/exon structure of the gene. Red and teal arrows indicate positions targeted by guide RNAs, which after break repair should produce an Exon 1-6 fusion allele that can be detected in genomic DNA using a 5’ primer at Exon 1 and a 3’ primer at Exon 6. **B.** Analytical PCR of indicated monoclonal human U251-MG cell CRISPR-Cas9 clones tested for IPMK-knockout (IKO). PCR reactions of Exon 1 using primers 1&2 (left lanes) show clone 2-F1 has no PCR product, PCR of Exon 6 using Primers 3&4 (middle lanes) shows PCR products in all lanes, suggesting the entire *IPMK* locus was present in the genomes of all clones tested, and PCR of “Exon 1-6” using Primers 1&4 (right lanes) shows presence the Exon1-6 fusion allele resulting from repair of Cas9-induced DNA breaks to exclude exons 2,3,4 and 5 of *IPMK,* thus clone 2-F1 is a validated genetic knockout of *IPMK*. Since these PCR reactions worked as expected, clones containing loss of the *IPMK* exon 1 locus with the presence of a PCR product corresponding to a fusion allele(s) or clear insertion/deletion (in/del) repair events in exon 1 were chosen for western analysis for the presence of IPMK protein. **C.** Western blots of whole cell lysates probed using antibodies directed against endogenous IPMK (upper) or actin (lower), suggesting indicated monoclonal CRISPR cell lines contain no detectable full-length IPMK protein compared to wild-type U251-MG cells. Monoclonal U251 cell line 2-F1 was the clone used for complementation with wild-type and kinase-dead IPMK. These data demonstrate monoclonal cell line 2-F1 is a U251-MG cell line that lacks detectable expression of full length human IPMK at both the protein and genomic DNA levels. This cell line was also confirmed to lack detectable IPMK enzyme activity by HPLC of ^3^H-metabolic labeling of inositol phosphates.

## Supplementary Figure S2

**Figure S2.**
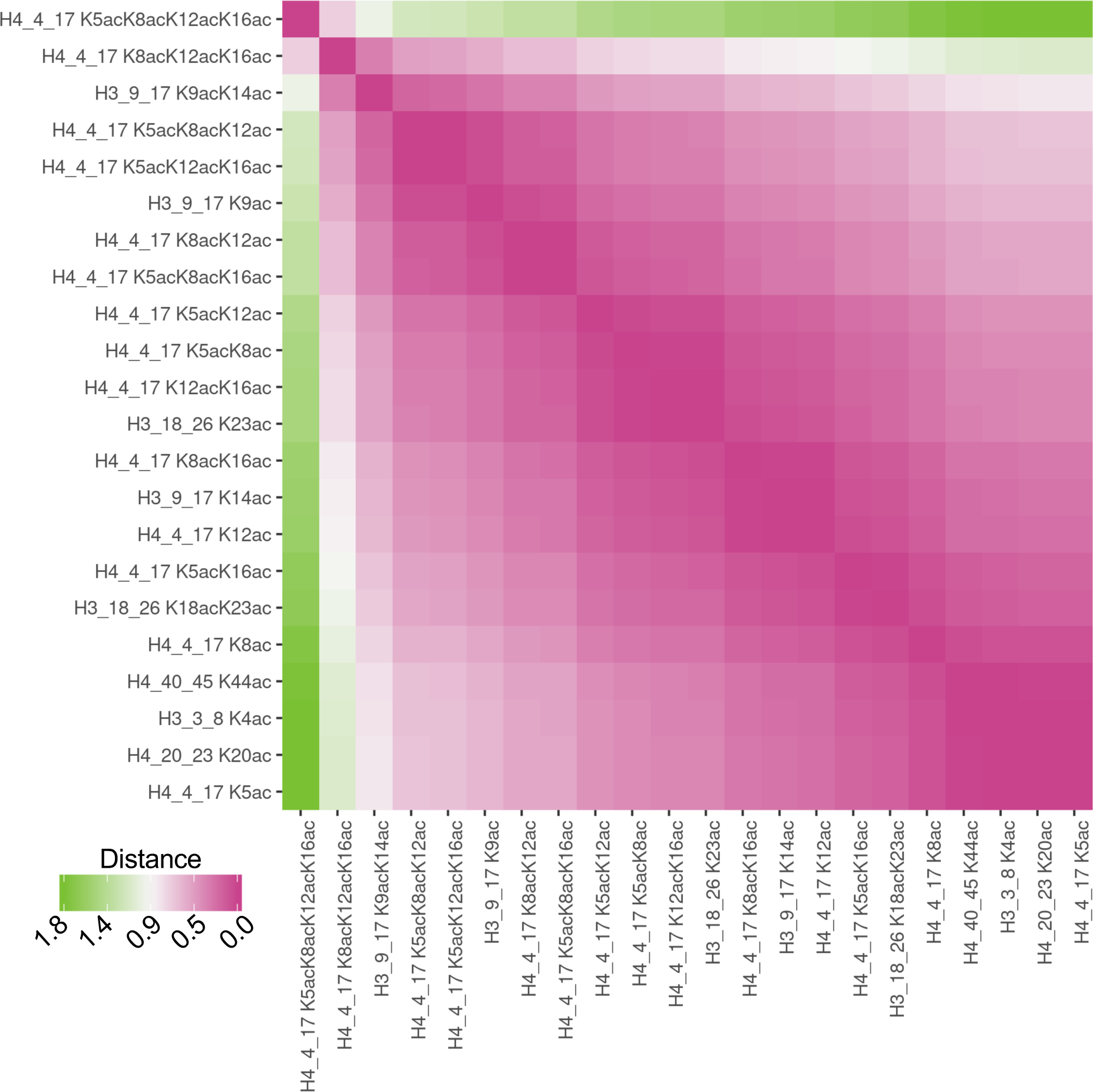
Complete labeling for pair distances of log2 fold changes from all acetylated peptides (IKO vs. WT) from proteomic mass spectrometry of Histone H3 and H4, showing the largest relative difference is for histone H4-K5acK8acK12acK16ac. Pair-distances of the log2 fold changes of all peptides with increased log2 fold changes in IKO vs. WT cells, only peptides which bear acetylation modifications were analyzed (phosphorylated and methylated peptides were excluded). These data suggest that the vast majority of acetylated peptides increase similarly (purple color ∼0 distance between most of the peptide pairs), however the most different peptides were the peptides that were most highly acetylated on histone H4, as detectable by proteomic histone mass spectrometry.

## Supplementary Figure S3

**Figure S3.**
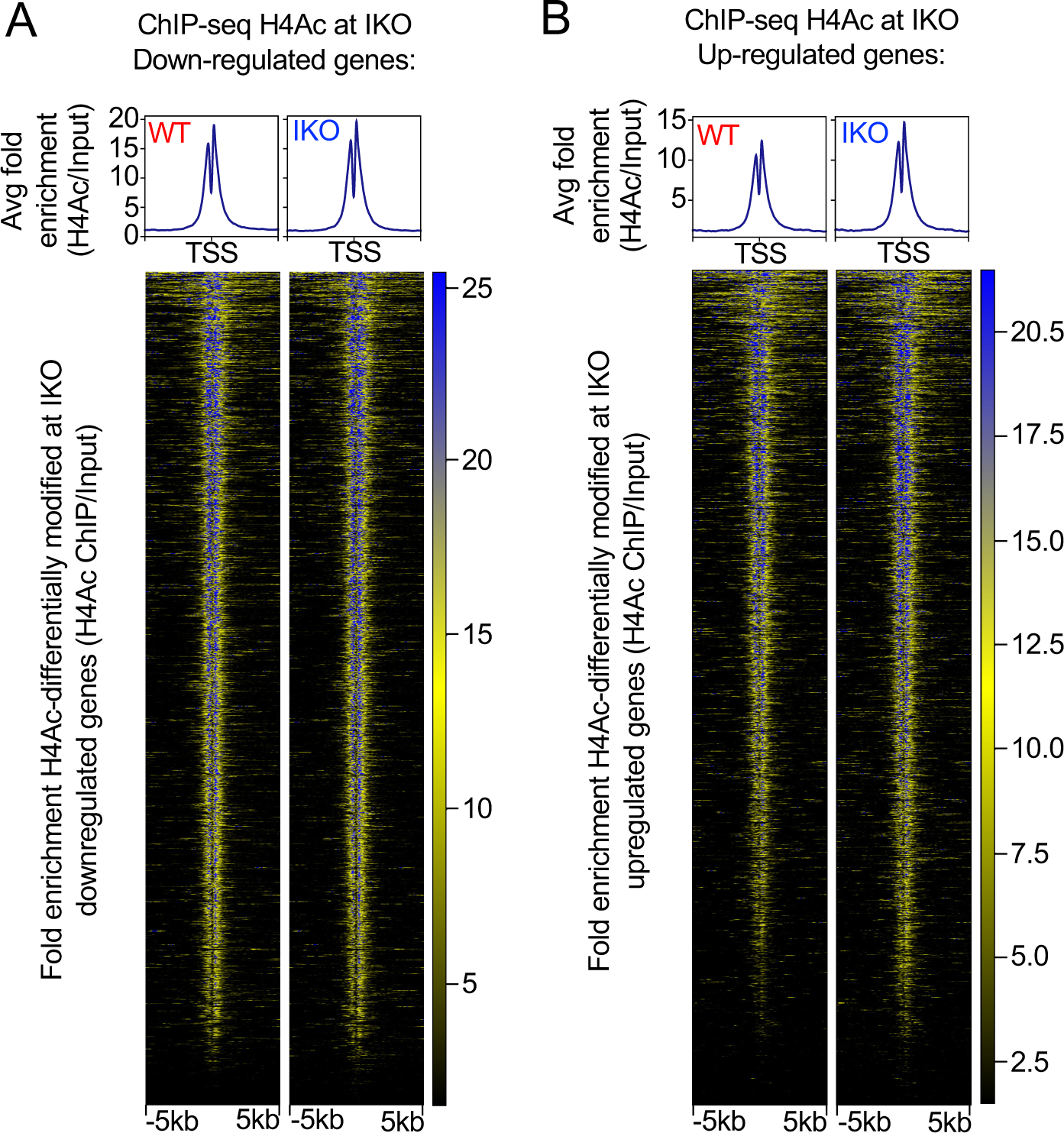
Histone H4Ac is enriched at IKO upregulated genes, but H4Ac is unchanged at IKO downregulated genes. **A.** Upper panel, average fold enrichment of ChIP against histone H4 poly-acetylation (H4Ac) from wild type (WT) cell chromatin (left) *vs*. IPMK-knockout (IKO) cell chromatin (right), aligned at the transcriptional start sites of all genes whose transcripts were **downregulated** by IPMK-knockout. Lower panel, heatmap of individual peaks at all genes downregulated by IPMK-knockout. **B.** Upper panel, average fold enrichment of ChIP against histone H4 poly-acetylation (H4Ac) from WT cell chromatin (left) *vs*. IKO cell chromatin (right), aligned at the transcriptional start sites of all genes whose transcripts were **upregulated** by IPMK-knockout. Lower panel, heatmap of individual H4Ac peaks at all genes upregulated by IPMK-knockout compared to wild-type cells. These data suggest histone H4 acetylation is not significantly changed at the transcriptional start sites of transcripts that were *down-*regulated by IKO, but H4 acetylation increases at start sites of transcripts that were *up*-regulated by IKO. These results suggest IPMK controls transcript abundance by two mechanisms, one of which associates with altered H4Ac and one that does not associate with any changes to H4Ac.

## Supplementary Figure S4

**Figure S4.**
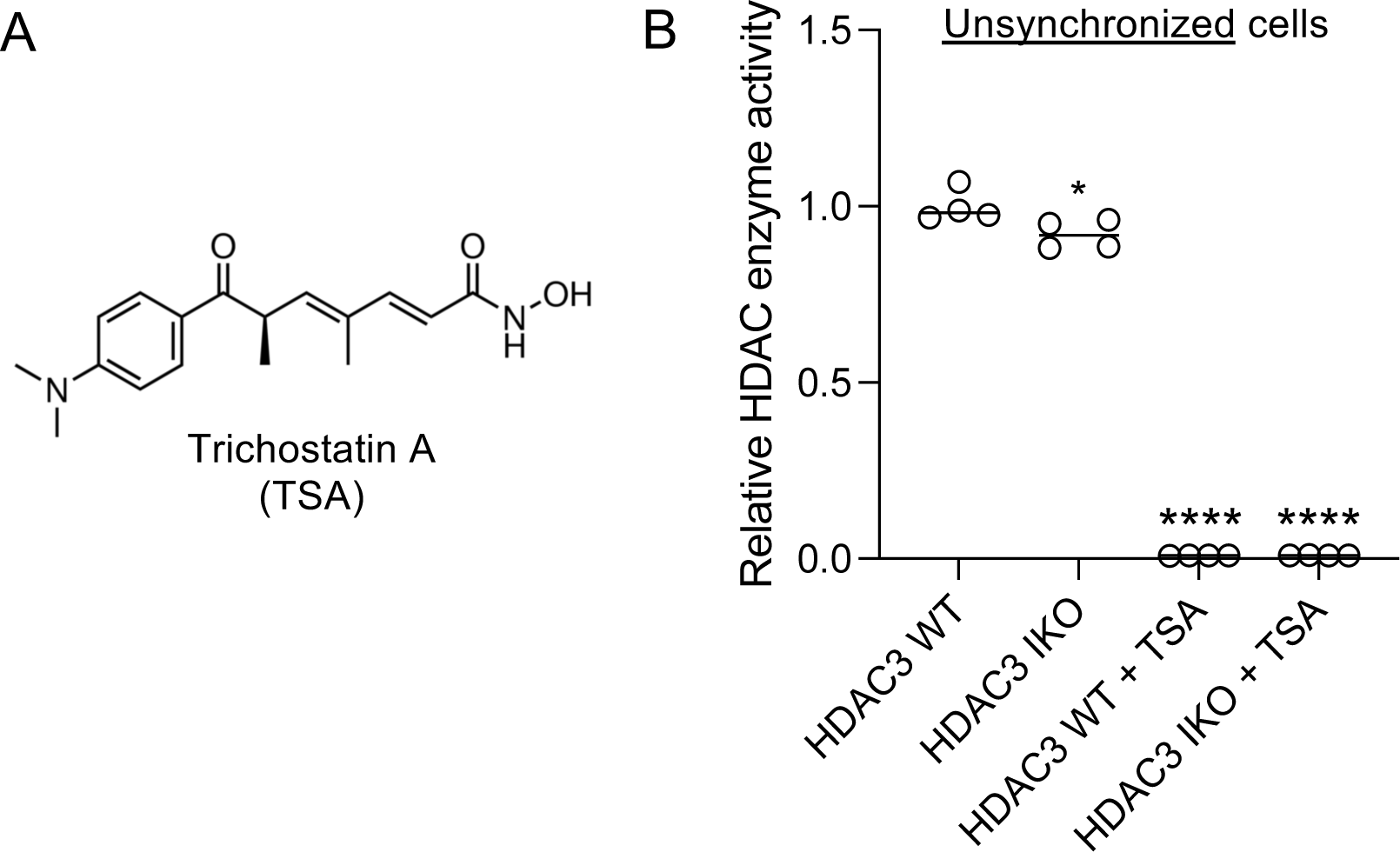
Trichostatin A reduces HDAC3 enzyme assay signal to background levels. **A.** Chemical structure of Trichostatin A, a well characterized inhibitor of HDACs. **B.** Relative HDAC deacetylase enzyme activity contained in HDAC3 immunoprecipitates from native whole cell extracts from unsynchronized cells, with or without the addition of Trichostatin A (Sigma), \**p=*0.0437, \*\*\*\**p*<0.0001 by unpaired t-test, signal from the Trichostatin A treated samples was reduced to background levels (empty wells in the assay plate), suggesting all the activity we measured in the HDAC3 immunoprecipitates was sensitive to Trichostatin A. The rather small effect size in the IKO HDAC3 immunoprecipitates on the left (not treated with Trichostatin A) is most likely due to these cells not being synchronized for these Trichostatin A control experiments.

## Supplementary Figure S5

**Figure S5.**
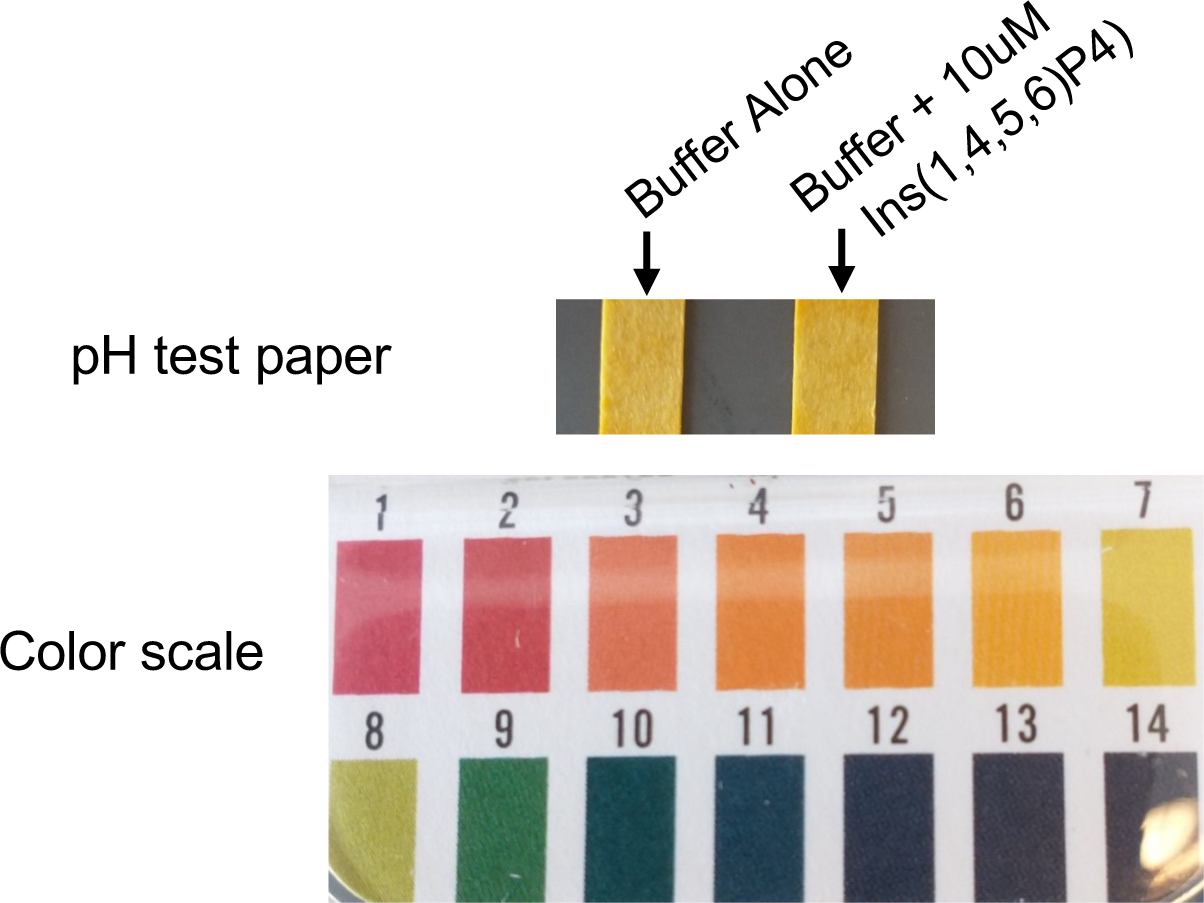
10uM *Ins(1,4,5,6)P4 does significantly change the pH of the HDAC enzyme reaction buffer.* IP4 contains 4 acidic phosphates that are known to acidify weakly buffered solutions, which could alter the results of the HDAC enzyme assay. To test if addition of Ins(1,4,5,6)P4 altered the pH of the HDAC enzyme reaction buffer, the reactions with or without Ins(1,4,5,6)P4 were spotted on pH test paper, which showed no change in pH upon addition of IP4, and the expected pH close to 7 as estimated from the pH strips. These data suggest addition of IP4 to the HDAC enzyme reactions does not alter the pH of the reactions.

## Supplementary Figure S6

**Figure S6.**
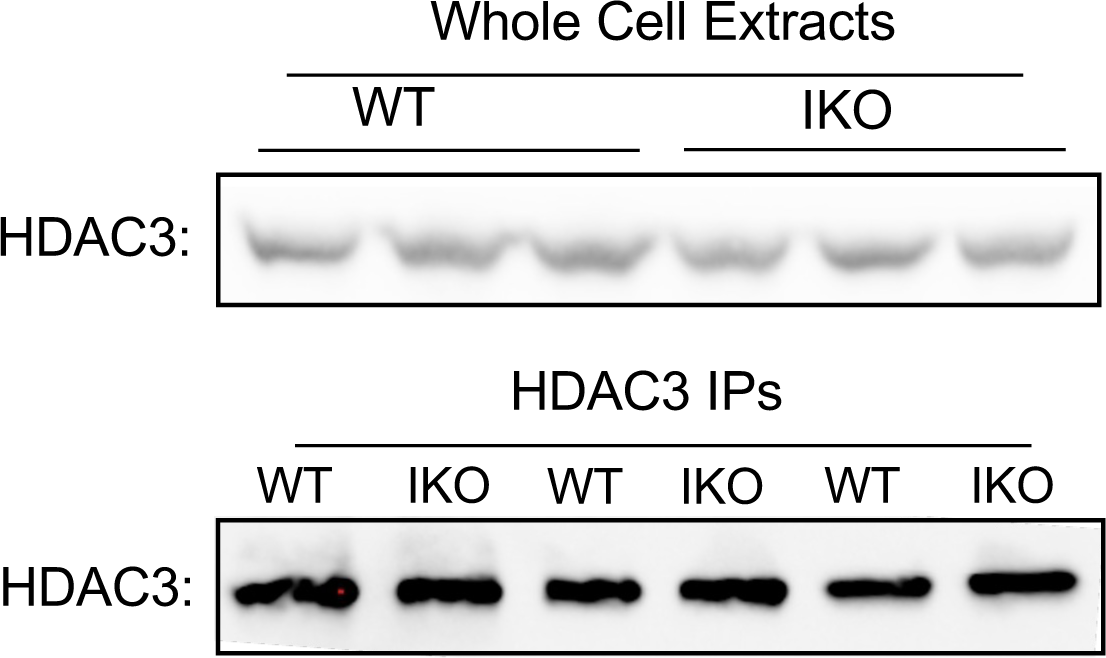
HDAC3 protein levels do not change in IKO cells. **A.** Western blots of HDAC3 from whole cell extracts (upper panel) and immunoprecipitated HDAC3 (lower panel) from three biological replicates, showing no change in protein levels in WT or IPMK KO (IKO) cells. These data suggest HDAC3 protein levels do not change upon IPMK knockout in U251-MG cells, and that immunoprecipitation pulled down equal amounts of HDAC3 from WT and IKO cells.

## Supplementary Figure S7

**Figure S7.**
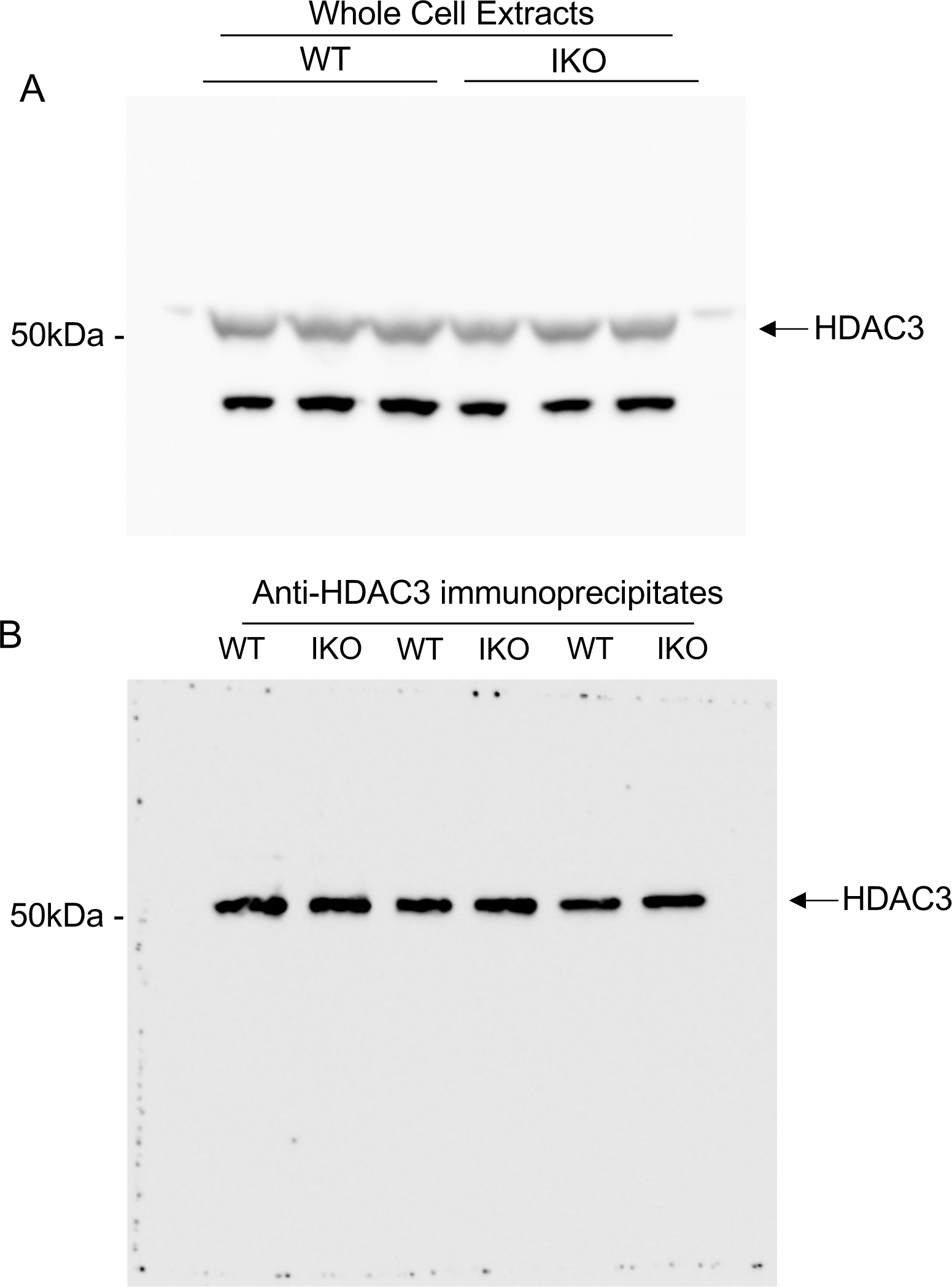
Whole western exposures showing HDAC3 protein levels do not change in IKO cells. **A.** Western blot of HDAC3 from whole cell extracts and **B.** immunoprecipitated HDAC3 from three biological replicates, showing no change in protein levels in WT or IPMK KO (IKO) cells in any of these samples, consistent with transcriptome analysis that showed no change to HDAC3 transcript abundance. These data suggest HDAC3 protein levels do not change upon IPMK knockout in U251-MG cells, and that immunoprecipitation pulled down equal levels of HDAC3 from the WT and IKO cells.

## Notes

### Competing Interest Statement

The authors have declared no competing interest.

