## Supplemental data for "IPMK regulates HDAC3 activity and histone H4 acetylation in human cells"

### Supplementary Figure S1

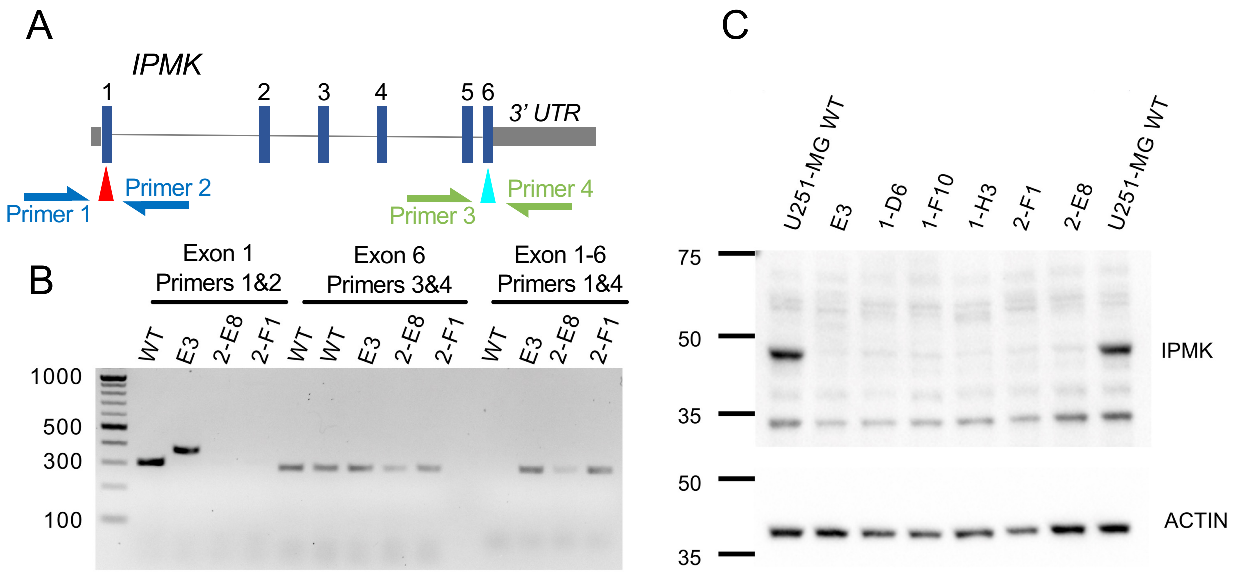


**Figure S1**. *Creation of IPMK knockout cells with CRISPR.* **A.** Model of human *IPMK* genetic locus, showing the intron/exon structure of the gene. Red and teal arrows indicate positions targeted by guide RNAs, which after break repair should produce an Exon 1-6 fusion allele that can be detected in genomic DNA using a 5' primer at Exon 1 and a 3' primer at Exon 6. **B.** Analytical PCR of indicated monoclonal human U251-MG cell CRISPR-Cas9 clones tested for IPMK-knockout (IKO). PCR reactions of Exon 1 using primers 1&2 (left lanes) show clone 2-F1 has no PCR product, PCR of Exon 6 using Primers 3&4 (middle lanes) shows PCR products in all lanes, suggesting the entire *IPMK* locus was present in the genomes of all clones tested, and PCR of "Exon 1-6" using Primers 1&4 (right lanes) shows presence the Exon1-6 fusion allele resulting from repair of Cas9-induced DNA breaks to exclude exons 2,3,4 and 5 of *IPMK,* thus clone 2-F1 is a validated genetic knockout of *IPMK*. Since these PCR reactions worked as expected, clones containing loss of the *IPMK* exon 1 locus with the presence of a PCR product corresponding to a fusion allele(s) or clear insertion/deletion (in/del) repair events in exon 1 were chosen for western analysis for the presence of IPMK protein. **C.** Western blots of whole cell lysates probed using antibodies directed against endogenous IPMK (upper) or actin (lower), suggesting indicated monoclonal CRISPR cell lines contain no detectable full-length IPMK protein compared to wild-type U251-MG cells. Monoclonal U251 cell line 2-F1 was the clone used for complementation with wild-type and kinase-dead IPMK. These data demonstrate monoclonal cell line 2-F1 is a U251-MG cell line that lacks detectable expression of full length human IPMK at both the protein and genomic DNA levels. This cell line was also confirmed to lack detectable IPMK enzyme activity by HPLC of $^{3}$H-metabolic labeling of inositol phosphates.

#### Supplementary Figure S2

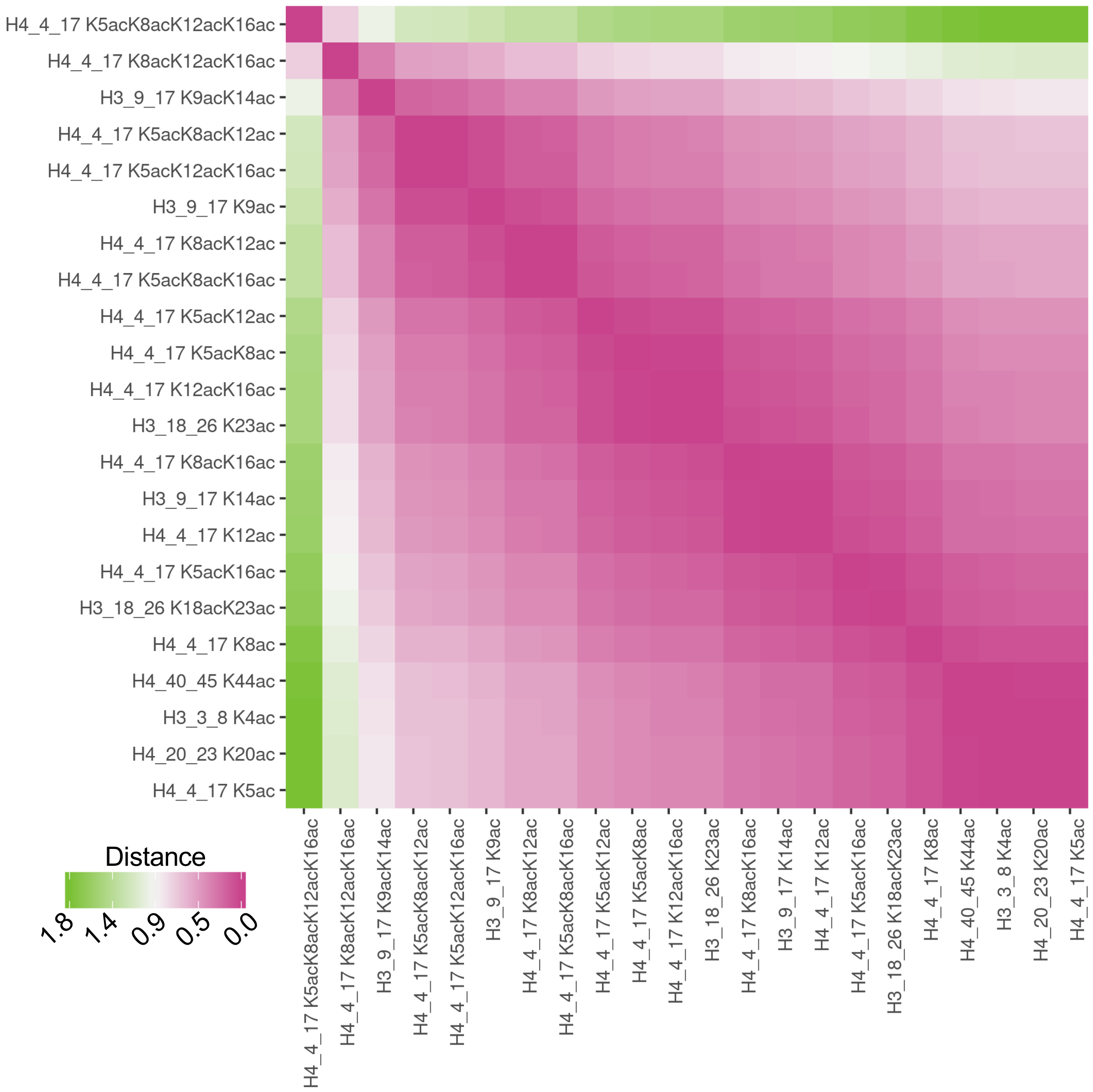


**Figure S2**. *Complete labeling for pair distances of log2 fold changes from all acetylated peptides (IKO vs. WT) from proteomic mass spectrometry of Histone H3 and H4, showing the largest relative difference is for histone H4-K5acK8acK12acK16ac*. Pair-distances of the log2 fold changes of all peptides with increased log2 fold changes in IKO vs. WT cells, only peptides which bear acetylation modifications were analyzed (phosphorylated and methylated peptides were excluded). These data suggest that the vast majority of acetylated peptides increase similarly (purple color ~0 distance between most of the peptide pairs), however the most different peptides were the peptides that were most highly acetylated on histone H4, as detectable by proteomic histone mass spectrometry.

#### Supplementary Figure S3

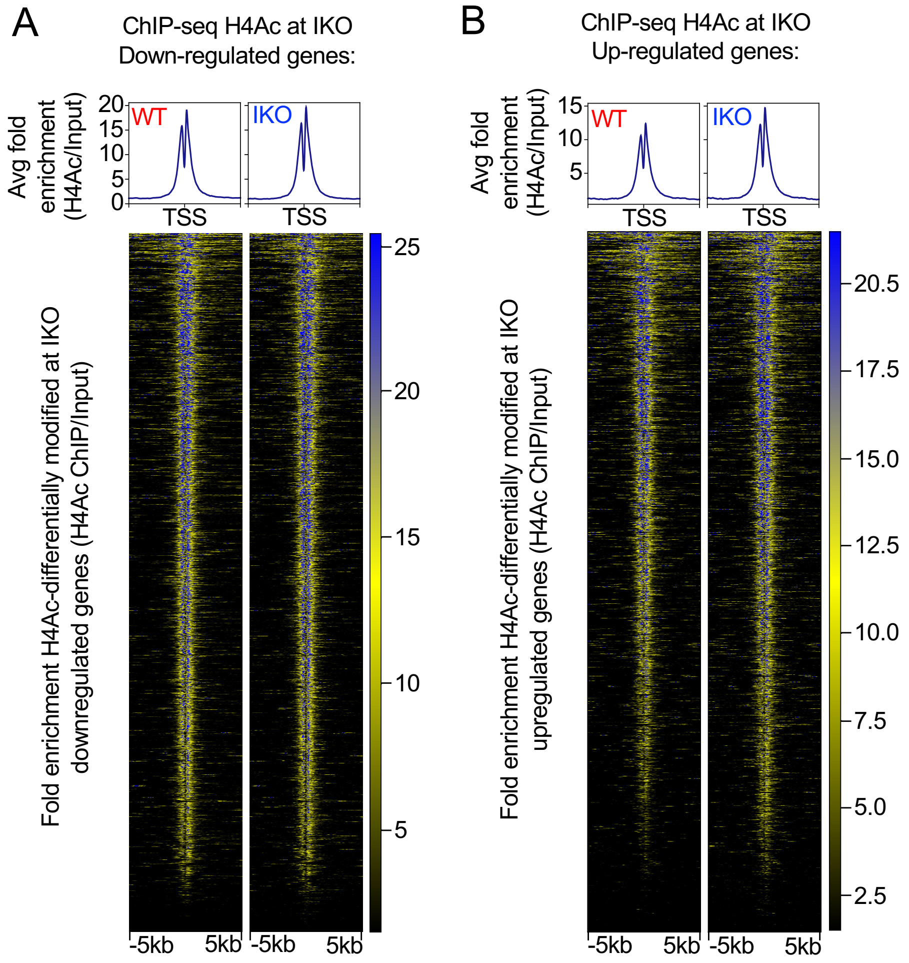


**Figure S3**. *Histone H4Ac is enriched at IKO upregulated genes, but H4Ac is unchanged at IKO downregulated genes.* **A.** Upper panel, average fold enrichment of ChIP against histone H4 poly-acetylation (H4Ac) from wild type (WT*)* cell chromatin (left) *vs*. IPMK-knockout (IKO*)* cell chromatin (right), aligned at the transcriptional start sites of all genes whose transcripts were **downregulated** by IPMK-knockout. Lower panel, heatmap of individual peaks at all genes downregulated by IPMK-knockout. **B.** Upper panel, average fold enrichment of ChIP against histone H4 poly-acetylation (H4Ac) from WT cell chromatin (left) *vs.* IKO cell chromatin (right), aligned at the transcriptional start sites of all genes whose transcripts were **upregulated** by IPMK-knockout. Lower panel, heatmap of individual H4Ac peaks at all genes upregulated by IPMK-knockout compared to wild-type cells. These data suggest histone H4 acetylation is not significantly changed at the transcriptional start sites of transcripts that were *down*-regulated by IKO, but H4 acetylation increases at start sites of transcripts that were *up*-regulated by IKO. These results suggest IPMK controls transcript abundance by two mechanisms, one of which associates with altered H4Ac and one that does not associate with any changes to H4Ac.

### Supplementary Figure S4

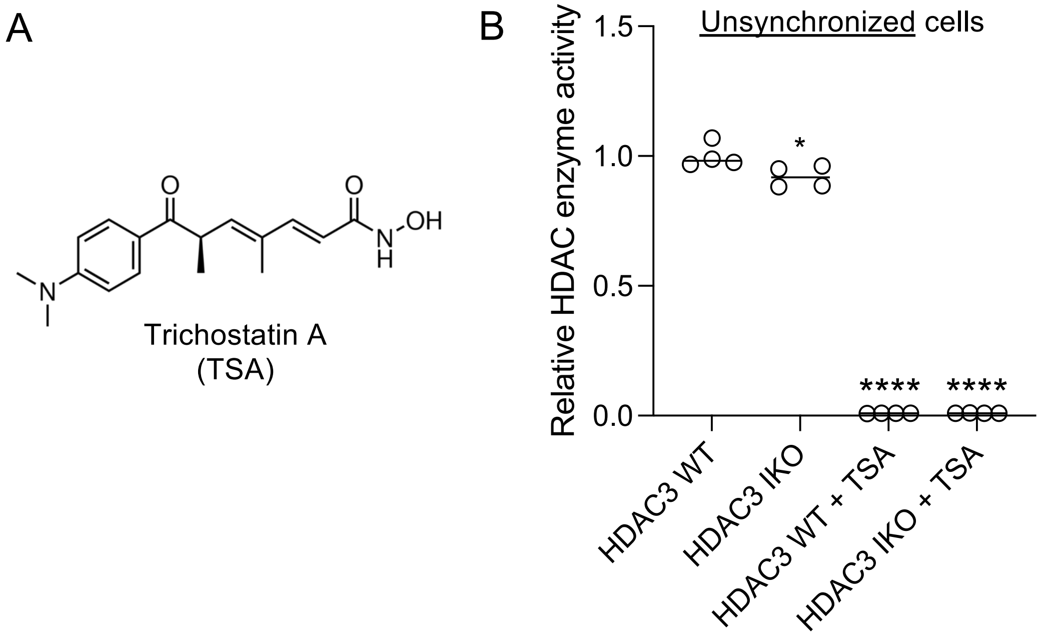


**Figure S4**. *Trichostatin A reduces HDAC3 enzyme assay signal to background levels.* **A.** Chemical structure of Trichostatin A, a well characterized inhibitor of HDACs. **B.** Relative HDAC deacetylase enzyme activity contained in HDAC3 immunoprecipitates from native whole cell extracts from unsynchronized cells, with or without the addition of Trichostatin A (Sigma), **p*=0.0437, *****p*<0.0001 by unpaired t-test, signal from the Trichostatin A treated samples was reduced to background levels (empty wells in the assay plate), suggesting all the activity we measured in the HDAC3 immunoprecipitates was sensitive to Trichostatin A. The rather small effect size in the IKO HDAC3 immunoprecipitates on the left (not treated with Trichostatin A) is most likely due to these cells not being synchronized for these Trichostatin A control experiments.

#### Supplementary Figure S5

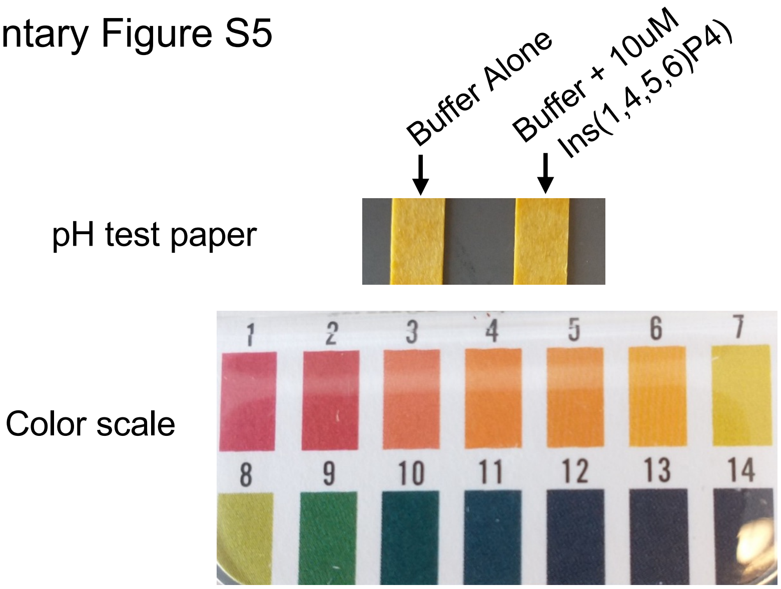


**Figure S5**. 10uM *Ins(1,4,5,6)P4 does significantly change the pH of the HDAC enzyme reaction buffer.* IP4 contains 4 acidic phosphates that are known to acidify weakly buffered solutions, which could alter the results of the HDAC enzyme assay. To test if addition of Ins(1,4,5,6)P4 altered the pH of the HDAC enzyme reaction buffer, the reactions with or without Ins(1,4,5,6)P4 were spotted on pH test paper, which showed no change in pH upon addition of IP4, and the expected pH close to 7 as estimated from the pH strips. These data suggest addition of IP4 to the HDAC enzyme reactions does not alter the pH of the reactions.

#### Supplementary Figure S6

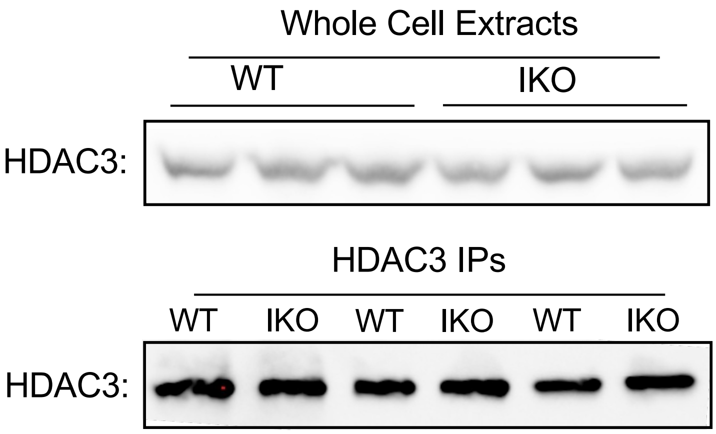


**Figure S6**. *HDAC3 protein levels do not change in IKO cells.* **A.** Western blots of HDAC3 from whole cell extracts (upper panel) and immunoprecipitated HDAC3 (lower panel) from three biological replicates, showing no change in protein levels in WT or IPMK KO (IKO) cells. These data suggest HDAC3 protein levels do not change upon IPMK knockout in U251-MG cells, and that immunoprecipitation pulled down equal amounts of HDAC3 from WT and IKO cells.

Supplementary Figure S7

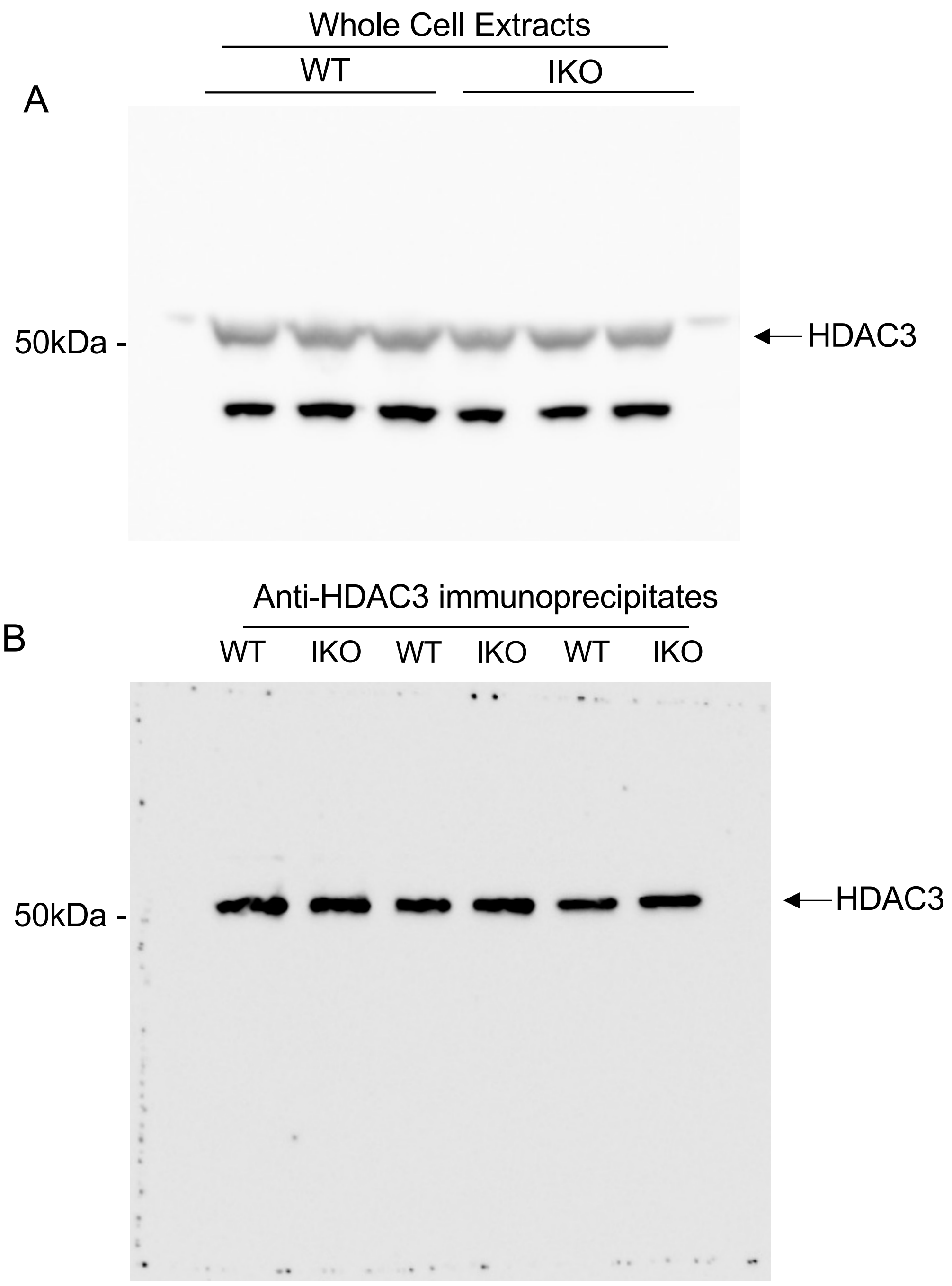


**Figure S7**. *Whole western exposures showing HDAC3 protein levels do not change in IKO cells.* **A.** Western blot of HDAC3 from whole cell extracts and **B.** immunoprecipitated HDAC3 from three biological replicates, showing no change in protein levels in WT or IPMK KO (IKO) cells in any of these samples, consistent with transcriptome analysis that showed no change to HDAC3 transcript abundance. These data suggest HDAC3 protein levels do not change upon IPMK knockout in U251-MG cells, and that immunoprecipitation pulled down equal levels of HDAC3 from the WT and IKO cells.
